# Geometric scaling of non-consumptive interactions generates sublinear density dependence and reshapes coexistence

**DOI:** 10.64898/2026.08.30.748073

**Authors:** Gaurav Baruah, KC Yogesh Kumar

## Abstract

The shape of density-dependence governs species persistence, and ecosystem stability. Yet, whether per-capita growth declines sublinearily, or superlinearily with density remains hotly debated. Growth rates across the tree of life have been shown to decline sublinearly with density, whereas theory founded on resource competition predicts the opposite. Here, we resolve this discrepancy and show that sublinearity can readily emerge from geometric constraints on consumer interactions. By linking inter-individual spacing, movement and interference rates, we derive two limiting-interference regimes, one of which “the well-mixed limit” recovers the form of classic Beddington–DeAngelis interference response. We then developed an individual-based model from first principles which reproduces the derived sublinearity response, and further use empirical data from published consumer-resource experiments that also bears the signature of sublinear density-dependence. Further, embedding the interference mechanisms underlying the emergence of sublinear density-dependence in coexistence theory opens a new regime for species coexistence where classical theory fails to predict. Our framework indicates that non-consumptive interactions are not merely a correction to resource competition but might be a distinct axis along which diverse communities may potentially coexist.

## 1 Introduction

Understanding how diverse species coexist remains one of the most fundamental questions in ecology (1). Dynamical models of species coexistence (2; 3; 4; 5; 6; 7; 8), or any other model that includes species interactions (9; 10; 11; 12; 13), most generally assume a linear density-dependent population growth. Density-dependence growth is a fundamental principle that shapes how species regulate themselves which has striking consequences on maintenance of biodiversity (14; 15). Classical phenomenological models of density-dependence such as the classic logistic growth rate, have helped in understanding and evaluating the mechanisms that maintain biodiversity (6; 16; 17). However, recent research has reported that density dependence is infact “sublinear” in nature, and is shown to be observed across myriad taxa(14; 15). Unlike the classic logistic growth model, where per-capita growth rates declines linearly with density, sublinear density dependence exhibits a convex, non-linear decelerating relationship (18; 19; 20; 21; 22; 23). Under sublinear density-dependence, population self-regulation (or intraspecific competition) is intense at low abundances, and then weakens as the population density increases. This pattern of decelerating density dependence has been supposedly observed across thousands of taxonomic groups, including mammals, birds, fish, insects, and eukaryotic phytoplankton (15; 21). However, such a pattern that has been reported, has been hotly debated as being inconclusive, as observational timeseries analysis that was used could have been impacted by statistical issues (19; 22; 23; 24). Nevertheless, if and when such sublinear density dependence prevails, it has shown that such non-linear self-regulation has profound implications for ecological stability. When species exhibit sublinear growth, the classic complexity-stability paradox is reversed; and increasing species richness now enhances community stability (15).

However, despite this phenomenological observation, the mechanistic emergence of sublinear density dependence remains poorly understood and highly controversial (22; 23). Standard, resource-explicit models of resource competition, such as the Monod formulations, typically predict linear or saturating-dependent growth as resources gets depleted (21). Furthermore, standard phenomenological formulations of sublinearity are plagued by a serious mathematical problem (15; 23). In these models, the per-capita growth rate diverges to infinity at near-zero abundances, creating an unrealistic “savior” effect that prevents species from ever going extinct (23). When this infinite growth is artificially corrected to reflect finite growth at low abundances, the sublinear model recovers the expected competitive exclusion, collapsing diversity as richness increases. In parallel, recent empirical studies employing continuous culture systems have reported strictly superlinear dynamics, calling into question the universality of sublinearity (22; 25) and suggesting that previous time-series analyses may suffer from statistical artifacts (22; 24; 25). Reconciling these contradictory findings requires a bottom-up framework showing how sublinearity can mechanistically emerge without mathematical invalid properties. This has important consequences towards understanding the mechanisms that promote and maintain diversity.

Historically, multiple frameworks have been proposed to explain the mechanisms that maintain diversity (2; 6; 26; 27; 28; 29). Among them, niche-based theories proposed that species maintain diversity by partitioning resources and thereby reducing competitive overlap (6; 30; 31), whereas neutral theory assumed functional equivalence among species and attributed biodiversity maintenance primarily to stochastic drift and dispersal (26). Contemporary ecology has sought to reconcile these perspectives through a niche-neutral continuum, formalized most prominently by modern coexistence theory (MCT) (6; 32). In MCT, species coexistence requires stabilizing mechanisms to be sufficiently strong to overcome fitness differences. Despite its conceptual power, applying this deterministic framework to complex communities remains challenging, partly because species are commonly represented as homogeneous units (4). This assumption overlooks the widespread phenotypic variation observed among conspecific individuals (33; 34; 35; 36), which can substantially alter competitive dynamics (37; 38; 39). Although individual-level variation might promote coexistence by hiding species-level differences, mathematical models of pairwise competition frequently show that high intraspecific variation in resource-use traits can destabilize coexistence and accelerate competitive exclusion (7; 37; 39). Mechanisms such as higher-order interactions (2; 28) or asymmetric plasticity may buffer these effects within particular regions of parameter space, but intraspecific trait variation generally disrupts coexistence (37; 39). Reconciling its ubiquity in natural populations with its potentially destabilizing effects thus represents a central challenge for coexistence theory.

To bridge the above gaps, we present a novel framework showing how sublinearity can mechanistically emerge from simple consumer-resource equations. We start from simple consumer-resource equations and introduce a non-consumptive interaction term that emerges by considering a geometric approach that is central to accessibility of resources under conspecific interactions. From this geometric approach, our conspecific interaction mechanisms leads to the emergence of two interference regimes: one which we call the “nearest neighbour mechanism” and the other we call the “well-mixed mechanism” that leads to the classic Beddington-DeAngelis interference form (40; 41; 42) that emerges from this approach. This geometric approach eliminates the low-density problems of phenomenological sublinear models (15; 23), and that while demonstrating that sublinearity naturally arises from very simple rules. To further evaluate the emergence of sublinear density-dependence from geometric constraints, we develop an individual based model of a population occupying a physically bounded environment. We devise this IBM from first principles, and show that from principles of spatial geometry and interference, sub-linearity can emerge as population density increases. Furthermore, we then use empirical FoRAGE (v5) (43) dataset to qualitatively test whether these mechanisms are detectable in published consumer-resource experiments, assembling interference series that differ in conspecific density. With this empirical support in hand, we then extend the framework to two competitive species and derive the MCT breakdown and show how prevalence of non-consumptive interference widens the coexistence region. Furthermore, we generalise this to trait-based competition of multispecies communities assembled with random interference matrices and evaluate how the structure of non-consumptive interference relative to resource competition determines species coexistence in presence of intraspecific variation. Our results show that non-consumptive interference can promote species coexistence despite high niche overlap enabling species to persist in the neutrality continuum. This geometric reformulation reconciles physical constraints with demographic models, providing a robust solution to the maintenance of biodiversity in complex communities.

### 2 Methods and Materials

### 2.1 Simple consumer-resource dynamics to sublinear density dependence

We begin with the simplest consumer-resource equations and then introduce a mechanistic derivation showing how sublinear density dependence can arise from non-consumptive interactions. In a well-mixed habitat, the dynamics of a single consumer, with density *N*, feeding on a single resource, with density *R*, can be written as (44; 45):

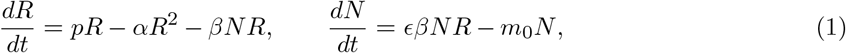

where *p* and *α* define the logistic growth dynamics of the resource, *β* is the consumer attack rate, *ϵ* is the conversion efficiency, and *m*_0_ is the per-capita mortality rate. Under the classical assumption that resource dynamics are fast relative to consumer dynamics, the resource remains near its positive quasi-equilibrium: 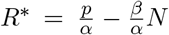. Substituting *R*^∗^ into the consumer equation leads to the familiar single-species logistic equation:

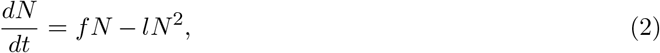

where, 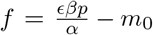 is the effective intrinsic growth rate and 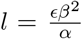 is the intraspecific competition coefficient. Eq. 2 exhibits strictly linear density dependence because the per-capita population growth rate 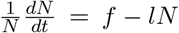, declines linearly with consumer density *N* . Next, we re-formulate the familiar consumer-resource equation by taking a geometric approach rooted in the physical accessibility of resources (Box 1, Box 2). Let us consider the same consumer-resource system, but assume that consumers occupy an enclosed habitat of area *A*, such as a two-dimensional feeding surface within a test tube or petridish. Consumers encounter and feed on resources, but during feeding they can also interfere with conspecifics. Their available time for feeding is thus partitioned between feeding and interference (Box 1 and Box 2). These interference effects come under the umbrella of non-consumptive interactions and could include aggressive interactions, avoidance behaviour, spatial aggregation, non-mating social interactions, or physical collisions. Although, these interactions do not directly impact consumers or resources, they potentially can reduce the amount of time that consumers can spend feeding. We thus assume that interference reduces the realised consumer attack rate to :

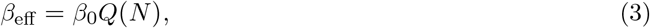

where *β*_0_ is the attack rate in the absence of interference and *Q*(*N*) ∈(0, 1] is the fraction of time spent feeding. Boxes 1 and 2 derive the relationship between inter-individual spacing, the rate of interference, and the fraction of time spent feeding. Box 1 presents two alternative mechanisms linking consumer density to interference, whereas Box 2 introduces the general density-scaling exponent *γ*. Substituting *β*_eff_ = *β*_0_*Q*(*N*) into the consumer-resource equations gives:

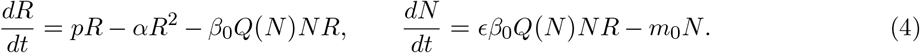

Applying the same timescale-separation argument, the resource remains near its positive quasi-equilibrium and substituting *R*^∗^(*N*) from the resource equation in 4, into the consumer equation yields:

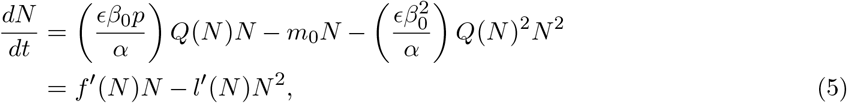

where 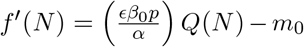 is the effective density-dependent population growth rate and 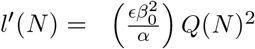 is the effective density-dependent intraspecific competition coefficient. The standard logistic or Lotka-Volterra dynamics are recovered when *Q*(*N*) = 1, corresponding to the absence of interference. When *Q*(*N*) *<* 1, non-consumptive interference reduces the realised attack rate and generates nonlinear density dependence. For 0 *< γ <* 1, the effect of density enters through the fractional power *N*^*γ*^, producing sublinear interference scaling (see Box 1 and Box 2). Crucially, the resource-dependent birth term is attenuated linearly by *Q*(*N*), whereas the intraspecific competition term is attenuated quadratically by *Q*(*N*)^2^. This structural asymmetry occurs because interference reduces both the focal consumer’s resource consumption and the amount of resource depletion generated by other consumers. We later show that this asymmetry has direct consequences for population persistence and coexistence. Unlike phenomenological formulations in which the per-capita population growth rate can diverge at low densities, the present model remains finite because lim_*N*→0_ *Q*(*N*) = 1. Consequently,

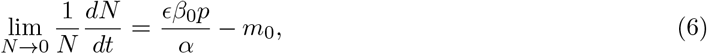

and the model returns smoothly to the standard consumer-resource dynamics when interference becomes negligible. Overall, what we show here is that a non-consumptive reduction in resource acquisition does not simply add another density-dependent mortality term. What it does is it simultaneously reduces consumer acquisition and resource depletion as well, thus changing the curvature and strength of demographic density dependence in a structurally asymmetric way. We show further this has meaningful significance in general to the debate of the presence of sublinearity density dependence.

#### Box 1

**From inter-individual spacing to interference**

Assuming, there are *N* consumers randomly distributed over an enclosed two-dimensional surface of area *A*. The consumer density is then given by 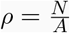. For a consumer, the mean area available would be given as *A*/*N* . Now, *A*/*N* has units of area. Next, we then quantify the characteristic linear spacing between neighbouring conspecific consumers. This scales as :

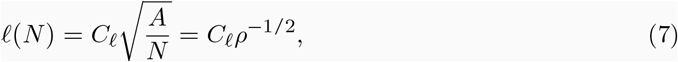

where *C*_*ℓ*_ is a dimensionless geometric constant. This constant could be dependent on the spatial distribution of consumers. One thing to note is that the spacing is not itself an encounter rate. To get to the encounter rate, we need to do the following. The rate of interference captures how individual conspecific consumers move through the space and how interference events happen. Let *λ*(*N*) denote the rate at which a focal consumer experiences interference. If the average uninterrupted feeding time is 1/*λ*(*N*), and each interference event causes an average feeding-time loss, say *τ*, then the fraction of time spent feeding is:

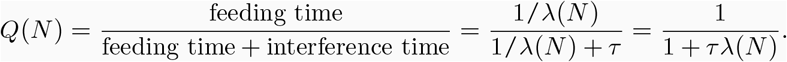

Here, we propose two alternative mechanisms that can connect consumer density to the rate of interference. One is what we call **Nearest-neighbour mechanism**: suppose that a focal consumer feeds until it moves across a distance comparable to the distance separating it from its nearest neighbour. This may occur when movement is persistent, and consumers are packed and aggregated, or when consumers respond directly to nearest conspecifics, or when they repeatedly cross their own neighbourhoods (Fig. 1A). If consumers move at an average speed *v*, the characteristic encounter time is 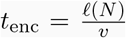. The interference rate is thus 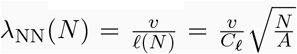.

**Figure 1.**
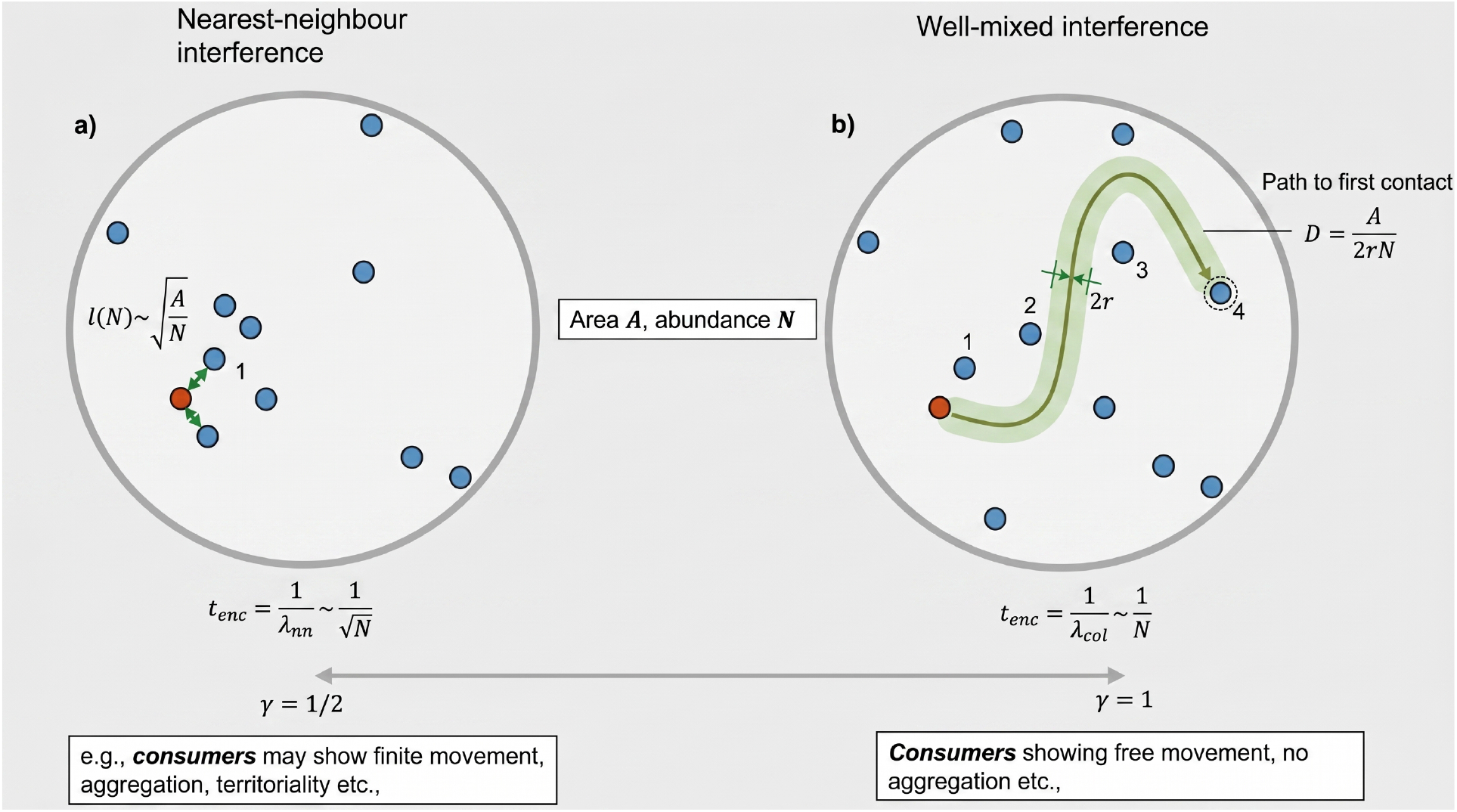
Non-consumptive interference generates sublinear density dependence under alternative encounter mechanisms. (**a**) *N* consumers occupy an enclosed habitat of area *A*, with characteristic inter-individual spacing 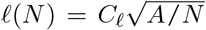. Under nearest-neighbour mechanism (**a**), a focal consumer (coloured orange) can cross a distance comparable (denoted by green arrows) to *ℓ*(*N*), giving an interference rate as *λ*_nn_ = *v*/*ℓ* ∝ *N* ^1/2^, where *v* is the movement speed. Under well-mixed collision (**b**), the focal consumer (coloured orange) moves in random directions and interferes when its path intersects the interaction area of radius *r* surrounding another consumer (it traverses pass individual 1, 2, and 3, and finally interferes individual 4, travelling a mean path *D* = *A*/(2*rN*) and giving *λ*_col_ ∝ *N* . Since path *D* ≥ *ℓ*, these mechanisms bracket the accessible range of density scaling. Under either mechanism, interference partitions the time budget: encounters at rate *λ* each cost a feeding-time loss *τ*,leaving a fraction *Q* = (1 + *τλ*)^−1^ for feeding. The resulting function *Q*(*N*) = (1 + *cN* ^*γ*^)^−1^ thus impacts density-dependence non-linearly and dependent on interference scaling factor, *γ*, with values implying spatial aggregation/attraction (*γ <* 1/2) or superlinear-like (*γ >>* 1) due to spatial avoidance behaviour.

Consequently,

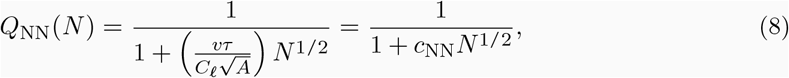

where 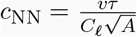 . The exponent *γ* = 1/2 arises when the time to interference is determined by the time required to traverse the characteristic nearest-neighbour distance in a habitat.

#### Box 2

**General interference scaling and the exponent γ**

**Well-mixed collision mechanism:** Alternatively, assuming that individuals could move in random directions, and in that case they could randomly interfere when their movement crosses with the interference distance of conspecifics. Particularly, the individual consumer might not necessarily traverse directly towards its nearest neighbour. Instead, it could pass through an area as it moves, and consequently interference could occur when its movement path intersects the interaction area surrounding another conspecific (Fig. 1B). In two dimensions, a consumer moving at average relative speed *v*_rel_ sweeps out an area per unit time proportional to 2*r*_int_*v*_rel_, where *r*_int_ is the interaction distance. If consumers occur at density *N* /*A*, the expected interference rate is therefore

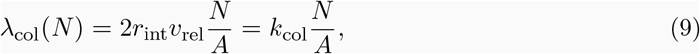

where *k*_col_ = 2*R*_int_*v*_rel_. Consequently, the well-mixed collision mechanism thus gives *γ* = 1 which has the form of the classic **Beddington–DeAngelis** interference curve. To be noted that our aim was not to derive the form of Beddington-DeAngelis curve. Instead we arrive at it from spatial geometry that included interference relationships. In addition, our geometric interference changes the relationship between intrinsic growth and resource-mediated competition in a way that Beddington-DeAngelis curve does not. Nearest-neighbour spacing may still scale as *N*^−1/2^ under this mechanism. This is because encounters depend on the number of interference areas crossed by a randomly moving consumer, rather than only on the distance to its nearest neighbour. Consumers in nature and their movements may lie anywhere between the nearest-neighbour interference mechanism (Box 1) and well-mixed collision mechanisms described here. For instance, consumers might show finite movement, or local avoidance or attraction, spatial aggregation, territoriality, or density-dependent changes in movement speed etc,. Consequently, the interference rate need not scale exactly as either *N* ^1/2^ or *N* . We thus represent the interference rate more generally as

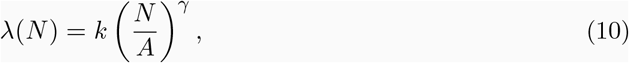

where *k* determines the overall magnitude of interference and *γ* determines how rapidly interference increases with consumer density. This exponent is not a phenomenological parameter, but arises from the geometry of consumer encounters. Substituting Eq. 10 into the feeding-time equation gives

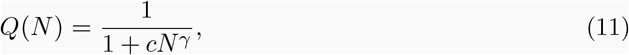

Where 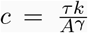. The exponent *γ* thus describes how consumer density is translated into nonconsumptive interference. More generally, values of *γ* describe a sublinear increase in the interference rate with consumer density, by taking the second derivative i.e.,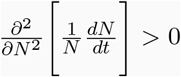, which is true for our *γ* = 1, 1/2. Values near 1/2 could arise if consumers avoid interfere with another in their neighbourhood, reduce their movement speed at high densities, or become spatially aggregated. Values very much greater than one could indicate a superlinear-like behaviour, potentially caused by avoidance behaviour, or individuals that are spatially segregated.

### 2.2 Individual based model and individual interference

We developed a spatially explicit individual-based model of a consumer species to investigate the emergence of sublinear density dependence from first principles of spatial geometry and interference. The IBM operates within a continuous 2-D circular arena, representing an environment where individuals can feed. We then distribute uniformly an initial population of consumers (*N*) in the confined area, each defined by a fixed body radius at the start of the simulation. The goal of this IBM was two fold: the first is to evaluate if conspecific interference within the arena can lead to the emergence of sublinear density dependence; the second goal was to evaluate whether our IBM can capture the interference scaling relationship *γ* = 1; 0.5 based on the two different regimes of interference mechanism, one that is random/well-mixed and the other being the nearest-neighbour interaction. Nearest-neighbour interaction can arise when consumers are attracted to each other in their own neighbourhood, depending on the individual’s affinity to aggregate with conspecifics. Thus, we use our IBM to evaluate three regimes of conspecific behaviour, one that is random and well-mixed (*ξ* = 0) where active individuals move through the arena via a random walk bounded by the arena’s physical perimeter. Here, *ξ* captures the individual behavioural interference towards conspecifics. The second is when individuals addition to moving around randomly, move towards and interfere (*ξ >* 0, nearest neighbour) or away from (*ξ <* 0) the nearest neighbour in their own interaction neighbourhood of some external radius (*R*_*interaction*_) and bump onto other conspecifics. The final third scenario is the null-model scenario where individuals move around the arena and there is essentially no interference that could occur. Crucially, this means that upon collision, both individuals that collide do not enter into any interrupted state for a predefined penalty duration. However, for the first two cases, individuals that collide enter into a temporary interrupted state for a predefined penalty duration. To mechanistically capture the effects of interference crowding, we calculate the pairwise Euclidean distances between all active individuals at every step. If the distance between any two individuals was less than twice their body radius, we registered a physical collision or interference. During this penalty period, individuals are immobilized and unable to consume resources. We had a feeding phase time loop representing loosely capturing a particular foraging season which gets impacted due to interference. However, feeding time does not get interrupted in the null-model case where conspecifics do not get immobilised due to interference.

Next, we translated these micro-scale spatial dynamics into a macro-scale population growth rate, which is governed by an individual energy-budgeting framework. During each micro-simulation loop, individuals that were active and successfully avoided collisions, accumulated energy by consuming a shared, finite resource pool at a constant feeding rate. As this resource depleted over time, individual feeding was limited by both unhindered access to space and resource competition. At the conclusion of the feeding phase, we subtracted a fixed metabolic maintenance cost from each individual’s total accumulated energy. Individuals that fail to meet this maintenance threshold die without reproducing. For those that survive, the remaining energy is multiplied by a conversion efficiency parameter to determine their total offspring. To rigorously quantify the shape of the resulting density dependence, we ran this IBM across an experimental density gradient, systematically varying the initial population size (*N*). By averaging the number of offspring produced per individual at each density level, we derived the realized per-capita growth rate, and also by quantifying the realised feeding efficiency, essentially *Q*, i.e., the time spent feeding divided by total time. Next, we estimate *γ* by fitting *Q*, which is the time spent feeding divided by total time, from our model results to 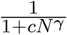 functional form. This approach enabled us to test whether spatial geometry, interference mechanisms (Box 1 and Box 2), and the resulting temporal non-consumptive interference inherently transform the classical linear intra-competition cost into a sublinear functional form, and further quantify the corresponding (*γ*) values across the three scenarios examined. For parameter values see table S2.

### 2.3 Empirical evaluation of interference-scaling mechanisms

We use FoRAGE version 5 database to test whether variation in feeding of consumers with increasing conspecific consumer density were qualitatively consistent with the geometric scaling relationship derived in Box1 and Box 2 (43). We restricted the analysis to only to consumer-resource interactions from the Forage database. Next, we compiled interference series by grouping observations that shared study, consumer and resource species, consumer life stage, foraging space, experimental duration, resource-replacement treatment and other reported experimental conditions, but differed in consumer number. If *P* consumers (predators) occupied an arena of area *A*, or volume *V*, we calculated the density of conspecifics potentially encountered by a focal consumer as *C* = (*P*− 1)/*A* or *C* = (*P*− 1)/*V*, respectively. We subtracted the focal consumer which ensured that a single-consumer treatment represented the absence of conspecific interference. We only analysed those data that contained at least three consumer-density levels. Consequently, the compiled dataset that we finally used contained 20 interference predator-resource series from 15 published sources from the FoRAGE database. We scaled resource density, consumer density and per-capita foraging rate by their respective within-series maxima. This was used to improve numerical stability for estimating the scaling interference parameter *γ*. To be noted that the FoRAGE analysis tests for the behavioural ingredient required by our mechanism, rather than directly testing and quantifying demographic sublinearity scaling values. Next, we used the following candidate models which followed directly from the theoretical accessibility function from Box 1 and Box 2, *Q*(*C*) = 1/(1 + *cC*^*γ*^). Substituting the effective attack rate *a*_eff_ = *a*_0_*Q*(*C*) which is impacted by the our theoretical *Q* function into a Type-II functional response gives:

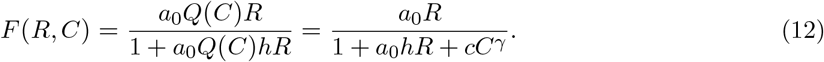

We thus compared a Type-II null model without interference,

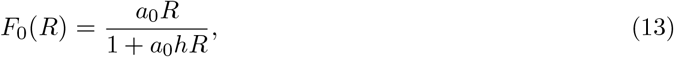

with two interference models differing only in the fixed value of *γ*. The first used the nearest-neighbour prediction *γ* = 1/2, whereas the second used the density-proportional, well-mixed prediction *γ* = 1 or the classic BD interference equation.

### 2.4 Model fitting and comparison

We fitted models independently to each interference of the predator-prey series using the original per-capita feeding-rate observations. We used nonlinear least squares with a Gaussian residual error model. For numerical fitting, we used Eq. 12 and was written as *F* (*R, C*) = *AR*/(1 + *BR* + *KC*^*γ*^), where *A, B* and *K* are non-negative fitted coefficients of the model. The null model estimated *A* and *B*. However, for each interference model, (i.e., *K*≠ 0), we had to additionally estimate *K*. Note, that since most FoRAGE data were not originally designed to estimate the scaling exponent and some did not replace consumed resources, we interpreted our statistical analysis as a qualitative comparison among candidate mechanisms rather than a quantitative estimation of the parameters. We fitted models fitted to the same observations and compared using the small-sample size corrected Akaike information criterion, AICc. For each consumer-resource series and model, we calculated ΔAICc relative to the model with the lowest AICc and obtained the Akaike weights. We considered models with ΔAICc *<* 1 as empirically equivalent under our criterion. The original values were retained when calculating Akaike weights and reporting model-selection tables. Also to be noted that when the *γ* = 1/2 and *γ* = 1 models were both within the equivalence threshold, we concluded that the series supported consumer interference but did not distinguish its density-scaling mechanism, as this was considered as qualitative support for our theory.

### 2.5 Two-species sublinear dynamics and modern coexistence theory

Our secondary goal was to scale the impacts of such interference into generalised coexistence mechanisms. We then extended our theoretical framework to generalised Lotka-Volterra equations to evaluate how our scaled interference mechanisms (nearest neighbour interference and well-mixed interference) can further impact species coexistence firstly in two species models, and then in larger multispecies competitive models. We extend Eq. 5 to two competing species, and that species *i*’s consumption is attenuated by its own interference factor *Q*_*i*_ while species *j*’s contribution to resource depletion is attenuated by *Q*_*j*_, the population dynamics are given as (see appendix section 1.2-1.5 for details):

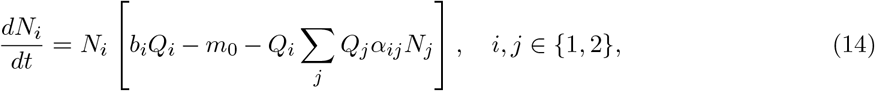

with the species-specific interference factor following from Box 1 and Box 2 and given generally as,

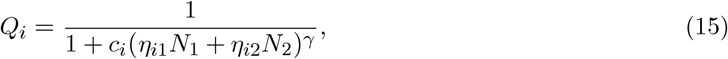

where *η*_*ij*_ is the contribution of species *j* to the interference experienced by species *i*. The matrix ***η*** thus captures the structure of non-consumptive interference, distinct from the resource-based competition matrix ***α***. When *c*_*i*_ → 0 or *η*_*ij*_ = 0 for all species, Eqs. 14-15 reduce to the two species generalised Lotka-Volterra competitive system.

#### 2.5.1 Invasion analysis, niche overlap and fitness ratios

We consider species *i* to be rare while species *j* is at its positive equilibrium, *K*_*j*_. The invader species then experiences interference only from the resident species, and the resident species’ depletion of resources is attenuated by its own interference factor. Based on this, the invasion criterion of species *i* becomes (see appendix for derivation):

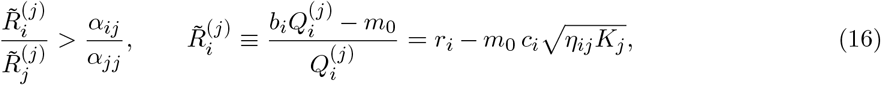

This equation is structurally similar to the Lotka-Volterra invasion criterion for two species, with the resident-conditional effective rates 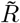 replacing the intrinsic rates *r* of Lotka-Volterra. Here *r*_*i*_ = *b*_*i*_ − *m*_0_. Next, we compared Eq. 16 with its Lotka-Volterra counterpart, and we see that interference facilitates the invasion of species *i* if and only if the invader species’ interference burden per unit intrinsic rate is smaller than the resident’s, i.e.

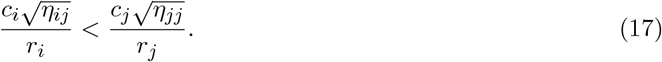

Following Chesson (6), we multiply the two mutual-invasibility inequalities by 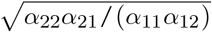 recover the MCT decomposition in the presence of interference. We find that the niche overlap 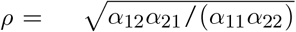 depends only on the competition matrix and is thus unaffected by non-consumptive interference. However, the fitness ratio, in contrast, becomes resident species-conditional given as:

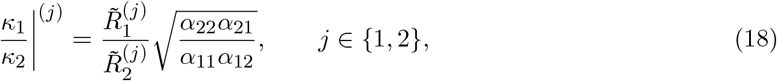

and mutual invasibility thus requires:

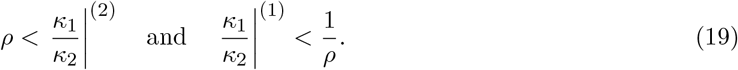

In the Lotka-Volterra limit 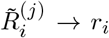 irrespective of the resident’s identity, and the two fitness ratios collapse to the single Chesson’s value. Under sub-linear density dependence they differ, and the resident-dependence of the fitness ratio is the analytical signature of interference in MCT: niche stabilisation operates exactly as in classical theory, while the fitness axis acquires structure that classical MCT does not capture. We further evaluate how the structure of the interference matrix *η* impacts species species coexistence. We quantify this by parametrising *η* around two ratios: the cross-species asymmetry *k*_asym_ = *η*_12_/*η*_21_ which captures whether species interferes with species strongly than reverse, the self-crowding asymmetry *k*_self_ = *η*_22_/*η*_11_, which quantifies whether species 2 experiences stronger self-interference than species 1. We then perform a two-dimensional sweep of each ratio against niche difference 1 − *ρ*, holding the other ratios fixed at unity, and evaluate species coexistence.

### 2.6 Trait-based consumer-resource eco-evolutionary dynamics

The species-level formulation does not capture the within-species phenotypic variation that mediates resource use. Next, we model phenotype-based competitive interactions that we again derive from McArthur’s consumer-resource equations. This formulation incorporates trait structure by reconciling quantitative genetics framework into Lotka-Volerra equations as followed from previous studies (38; 39). We assume that individuals of consumers are characterised by a quantitative phenotype, *z*, that determines how efficiently they consume resources of varying quality. Such a trait is in the quantitative genetic limit, and is thus assumed to be normally distributed. In such a case, the per-capita growth rate of a phenotype *z* for species *i* in the presence of interference with other consumer phenotypes can be writtenas (see full derivation in appendix section 2):

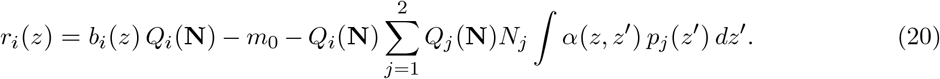

where *α*(*z*, − *z*^′^) = exp(−(*z* − *z*^′^)^2^/*w*^2^) is the trait-based competition kernel between phenotypes (38; 39); here we assume that non-consumptive interference factor crowding is trait-independent, i.e., the *η* matrix belonging to *Q*_*i*_(**N**), where the diagonal elements (*η*_*ii*_) of the matrix correspond to non-consumptive self-interference and off-diagonal elements correspond to non-consumptive interspecies interference that are driven by some unknown trait which we model from as a random variable drawn from a scaled random uniform distribution (see below).

Integrating Eq. 20 over the trait distribution *p*_*i*_(*z*) yields the species-level population dynamics which can be written as

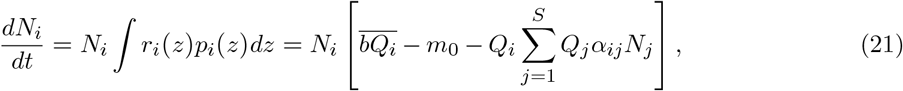

and evolutionary dynamics of the mean phenotype is given as

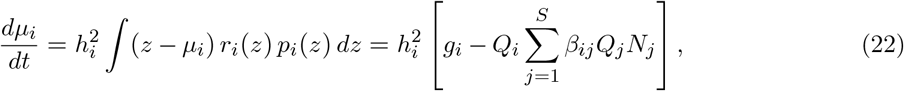

Here, the *α*_*ij*_ is the competition coefficient which captures on average how strongly species compete based on their mean phenotypic values. Similar species will compete strongly for resource use than dissimilar species, with *w* capturing the width of the competition kernel, *β*_*ij*_ captures how selection due to competition shapes the trajectories of mean species trait value; 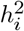 is the heritability of the focal trait, which is the fraction of phenotypic variance and genetic variance. The elements of the species-level interference matrix are drawn from random uniform distribution, which we will detail below. When *c*_*i*_ = 0 for all species, Eqs. 21 and 22 reduce exactly to the Lotka-Volterra eco-evolutionary model. Based on this, firstly we simulate eco-evolutionary dynamics for two species competition models under interference, and integrate MCT fitness ratio and niche overlap emerging from equation 21 and 22 for two species to demonstrate how intraspecific variation, eco-evolutionary dynamics and non-consumptive interference impacts eco-evolutionary dynamics of MCT and thus the coexistence region. In this case, two species trait-based niche overlap can be quantified from equations 21 and 22 (see appendix section 3 for derivation) as :

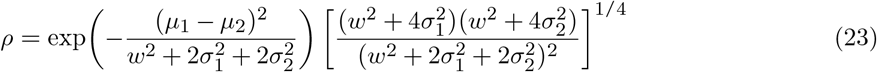

And, fitness ratio can be then quantified as:

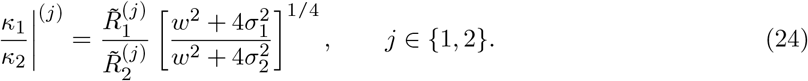

From this, the interference term *Q* directly impacts the fitness ratio through 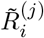, and we through simulations evaluate the impact interference scaling factor of *γ* ∈ [0.5, 1], and the interference matrix coefficient, i.e., ratio of 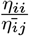 on niche overlap, and fitness ratio in the presence of high versus low intraspecific variation i.e., *σ*_*i*_, *σ*_*j*_.

After the example two-species competition model, we then generalise and extend our simulations to *S* species, where we vary initial *S* from 30, 40, 50 and evaluate how varying regimes of non-consumptive interference matrix impacted species coexistence at eco-evolutionary equilibria. Specifically,To examine how conspecific interference affects community persistence, we varied the diagonal elements of the interference matrix, ***η***, while holding its off-diagonal elements fixed within each simulated community. We first defined a community-specific interference scale using the mean intraspecific competition coefficient: 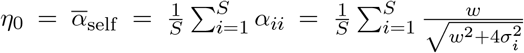. The off-diagonal coefficients, representing heterospecific interference, were drawn independently as: *η*_*ij*_ ∼ *η*_0_ U [0, 2]. Their expected value was therefore E [*η*_*ij*_] = *η*_0_, although their realised mean varied slightly among communities. We then set the diagonal coefficients, representing conspecific interference, to *η*_*ii*_ = *ψη*_0_. Thus, *ψ* measures the strength of conspecific interference relative to the mean strength of intraspecific resource competition and, in expectation, relative to the mean heterospecific interference coefficient. Values of *ψ <* 1 represent weaker conspecific than average heterospecific interference, *ψ* = 1 represents equal expected strengths, and *ψ >* 1 represents stronger conspecific interference. We varied *ψ* ∈ {0.2, 0.4, 0.8, 1, 2, 4, 8, 10 }. For each community, the off-diagonal interference coefficients were generated once and retained across all values of *ψ*, ensuring that only the strength of conspecific interference changed across the parameter sweep. Scaling ***η*** by *η*_0_ also kept the interference coefficients comparable in magnitude to the trait-based competition coefficients. We ran fully factorial simulations varying *ψ* ∈ [0.2, 0.4, 0.8, 1, 2, 4, 8, 10], crossed with *γ* ∈ [0.5, 1] and with low trait variance *σ* ≈ *U* [0.001, 0.005] versus high trait variation *σ* ≈ *U* [0.01, 0.1] each with fifty replicates, and compared our results to the case of generalised trait-based Lotka-Volterra model without interference, i.e., *c*_*i*_ = 0, for all species. As *ψ* values increase, intraspecific interference increases relative to interspecific interference for a particular community with *S* species. We then estimated inverse Simpson’s diversity index at eco-evolutionary equilibrium.

## 3 Results

### 3.1 Sublinear density dependence emerges from geometric constraints

We first verified whether the geometric interference mechanism (Box 1, Box 2, Fig. 1) produces sub-linear density dependence. We show that per-capita growth rate declines linearly as density increases as *c* = 0 which is the classic Lotka-Volterra model in figure 2A .When *c >* 0 and *γ* = 1 or *γ* = 0.5, the per-capita growth rate shows a concave decline as density increases characteristic of sublinear density dependence. The growth rate approaches zero more slowly at high density as shown in figure 2A. What we observe in figure 2A is the analytical signature of the *Q*^2^ attenuation of intraspecific competition in Eq. 5.

**Figure 2.**
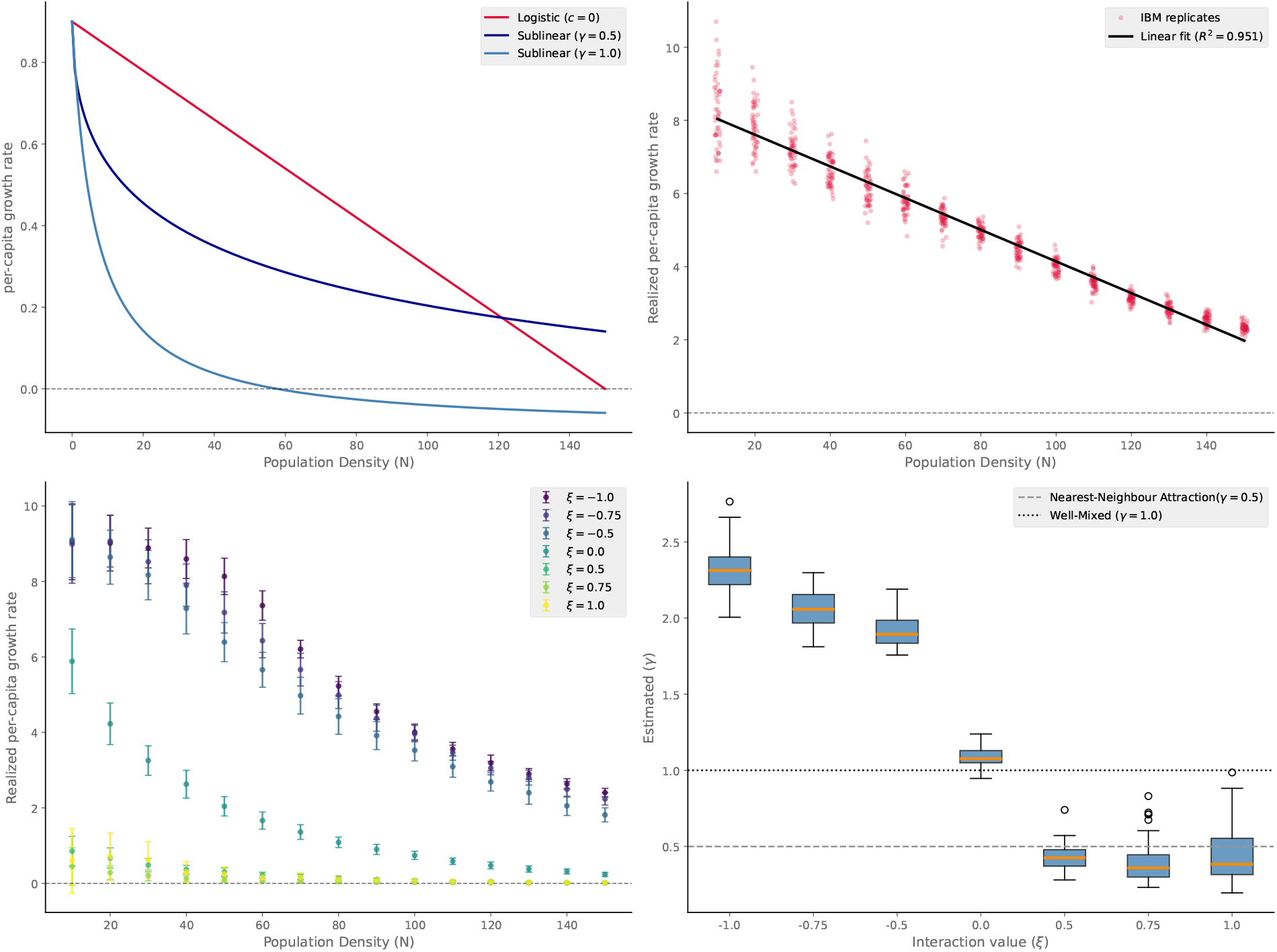
Emergence of density-dependent regimes. (A) Analytical per-capita growth rate plotted against equilbrium population density for logistic (crimson color, *c* = 0), and for sublinear models (*γ* = 0.5, *γ* = 1). Parameters used *m*_0_ = 0.1; *b* = 1; *c* = 0 for LV and for sublinear models parameters used *m*_0_ = 0.1; *b* = 1; *c* = 0.15. (B) realised per-capita growth rate from our IBM in relation to population density, where each point (*n* = 50) represent replicate simulations, and the line is a fitted linear model, with estimated *R*^2^ = 0.95. For (B): in our circular arena, individuals move via random walk and collisions do not produce any time-penalty. Thus, individuals can consume without any interference, this produces linear decrease in realised growth rate as density increases. (C) In the same IBM, we introduce nearest-neighbour interaction (*ξ*) in some neighbourhood(*R*_*interaction*_) and introduce a time penalty every time individuals collide and interfere. This produces a differential growth behaviour in relation to population density for varying interaction *ξ*. (D) Estimated *γ* from realised feeding efficiency *Q* transitions from a superlinear (*γ >* 1) to sublinear (*γ <* 1) growth regime as the nearest-neighbour interaction changes from repulsion (*ξ <* 0) to attraction (*ξ >* 0). Mean and Standard deviation of *γ* estimates for increasing *ξ*, [(3.08, 0.24), (2.88, 0.16), (2.57, 0.12), (1.11, 0.10), (0.42, 0.09), (0.40, 0.08), (0.301, 0.052), (0.37, 0.08)] respectively. Parameters used for this IBM are detailed in table S2 in appendix.

To confirm that the geometric derivation of *Q*(*N*) reflects a real mechanistic process rather than a phenomenological one, we implemented the IBM of consumers occupying an enclosed habitat, where feeding is interrupted upon conspecific encounter and a penalty time must elapse before feeding resumes. When the penalty time is set to zero (removing the interference cost, Fig. 2B), we see the dynamics revert toward the classical linear form. Diving deeper into this, we then evaluated three regimes of consumer behaviour given by *ξ*; *ξ* = 0, and *ξ >* 0 in the IBM characterises the well-mixed/random movement behaviour and nearest-neighbour interference or spatial aggregation towards conspecific behaviour respectively. We observed that realised per-capita growth rate falls sublinearily in relation to increases in density dependence (Fig. 2C) for both the *ξ* values. When *ξ <* 0, characteristic of consumers actively avoiding nearest-neighbours, we observed threshold-like superlinear density-dependence, but only when consumer density was low (Fig. 2C). When we fitted realised feeding time i.e., *Q* to different values of consumer behaviour quantified by *ξ*, our estimated *γ* from the simulations very well represented our nearest-neighbour interference mechanism and well-mixed random interference mechanism (Fig. 2D), although with some variability (Fig. S4).

### 3.2 Empirical evidence of sublinear density-dependence

Of the 20 interference series compiled from the FoRAGE database, 12 (60%) supported an interference model over the no-interference type-II null by more than two AICc units, and in these series the null model was frequently rejected decisively (Δ AICc up to 46.6; Fig.3A). Among the series supporting interference, both density-scaling mechanisms were represented: the nearest-neighbour prediction *γ* = 1/2 and *γ* = 1 i.e., the well-mixed interference mechanism (Fig. 3A-B). Taken together, these results indicate that consumer feeding rates in a substantial fraction of published functional-response experiments decline with conspecific density in a manner qualitatively consistent with the sublinear interference function *Q*(*C*). Akaike weights for the interference scaling mechanisms signify that both interference models can in some case be equally supported (Fig. 3B).

**Figure 3.**
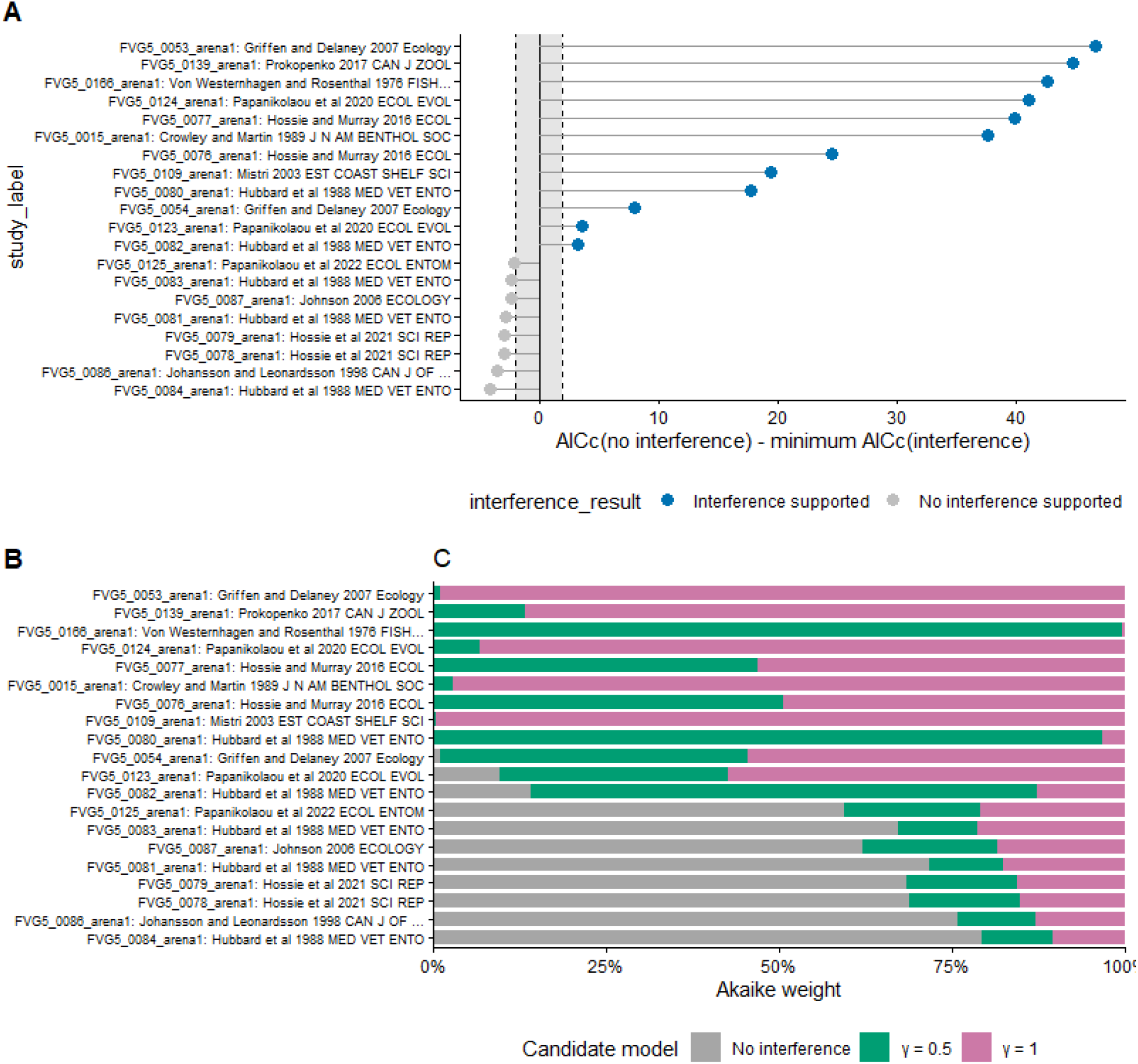
Empirical data from FoRAGE database shows sublinear density dependence can be present. A) Across *n* = 20 times series of consumer-resource feeding experiments, we found considerable support for the presence to sublinear density dependence within the regimes of *γ* = 0.5 and *γ* = 1. Positive values denote the presence of interference for studies on y-axis, and negative values indicate the absence of consumer interference. B) Horizontal bar stacked plots of Akaike weights providing weigth to interference-scaling mechanisms (green for *γ* = 0.5 and pink for *γ* = 1) in contrast to null-model with no interference (grey colour) for the studies on y-axis.

### 3.3 Sub-linearity and the structure of interference matrix widens the coexistence region

Under pure Lotka-Volterra dynamics competitive dynamics with *c* = 0, (Fig. 4A), coexistence requires the fitness ratio 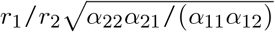 to lie between *ρ* and 1/*ρ*. The green coexistence region converges to a single point at zero niche difference (1 − *ρ* = 0). Two species coexistence at complete niche overlap requires exactly equal fitness ratios, and any demographic asymmetry excludes one species. Under the sublinear models *c >* 0, *γ* = 1/2, *γ* = 1,(Fig. 4B-C), the coexistence area widens at zero niche difference. We find that coexistence remains possible for fitness ratios in the interval as *ρ* → 1. Sub-linear density dependence emerging from interference thus promotes coexistence at high niche overlaps at which classical Lotka-Volterra dynamics predict competitive exclusion. This indicates that interference causes widening of the coexistence region due to the modified fitness axis as niche overlap *ρ* itself is unchanged by sub-linearity.

**Figure 4.**
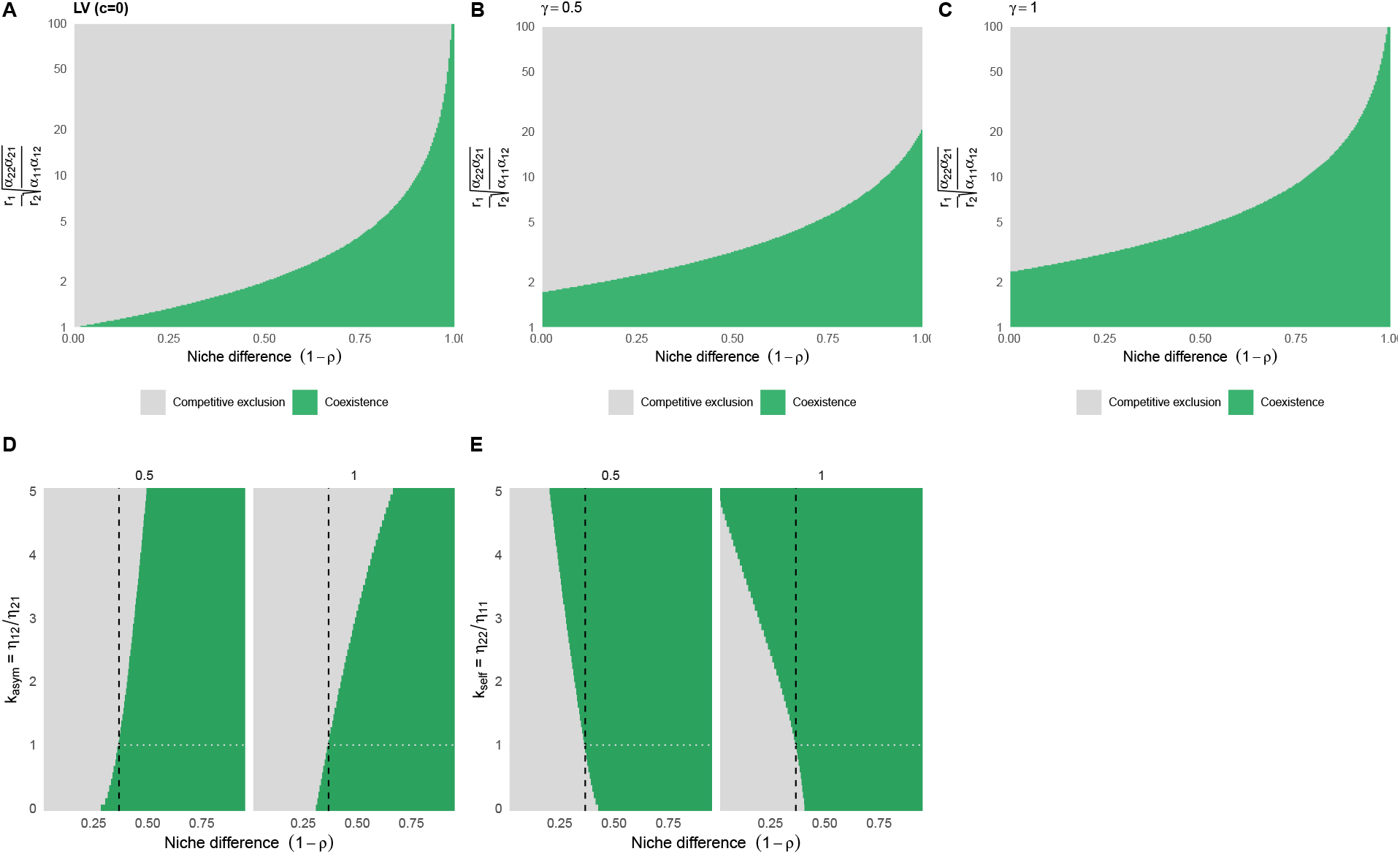
Non-consumptive interference expands coexistence parameter space for two-species. A-C: Coexistence regions in the MCT space of fitness ratio and niche difference. Green colour indicates parameter combinations satisfying mutual invasibility; grey indicates competitive exclusion. A) Generalised LV dynamics (*c* = 0), in which coexistence requires condition equation 1 to be valid, and the coexistence region vanishes when niche overlap approaches 1. B-C) With interference i.e., *γ* = 0.5 (nearest neighbour mechanism), and *γ* = 1 (well-mixed interference), the coexistence region retains finite space even at zero niche difference, so that species with very similar resource use can still coexist. (D-E) This coexistence under non-consumptive interference (*γ* = 0.5, *γ* = 1), depends further on the structure of the interference matrix *η*. Vertical dashed black lines demarcate Lotka-Volterra coexistence boundary (left of it LV predicts exclusion), and horizontal dotted grey lines mark symmetric interferences. D) Cross-species interference assymetry *k*_*asym*_; (E) Self-interference asymmetry, 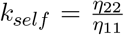. Green colour regions extending left of the dashed vertical line identify parameter spaces where coexistence is promoted with interference, and where resource pairwise competition however promotes exclusion. Grey colour indicates exclusion of species 1. Parameter values *α*_11_ = *α*_22_ = 1, *m*_0_ = 0.25; *c*_*i*_ = 1, where *c >* 0. For A-C, we assume *η*_*ii*_ = 1, *η*_*ij*_ = 0.01. For D: we varied (*k*_asym_ = *η*_12_/*η*_21_) from 0.01 to 5, while keeping (*η*_11_ = *η*_22_ = *η*_21_ = 0.5) fixed and setting (*η*_12_ = 0.5*k*_asym_). For E: we varied (*k*_self_ = *η*_22_/*η*_11_) over the same range,keeping (*η*_11_ = *η*_12_ = *η*_21_ = 0.5) fixed and setting (*η*_22_ = 0.5*k*_self_).

Although sublinearity *γ* = 1, *γ* = 1/2, generically widens the coexistence region in MCT space, whether it also rescues coexistence in a given system could directly depend on the structure of the interference matrix ***η***. We quantify this dependence by parametrising *η* around three biologically meaningful ratios: the cross-species asymmetry *k*_asym_ = *η*_12_/*η*_21_ (does species 2 interfere with species 1 more than vice-versa?), the self-crowding asymmetry *k*_self_ = *η*_22_/*η*_11_ (does species 2 self-interfere more than species 1?). We show that the two-dimensional sweep of each ratio against niche difference 1 − *ρ*, holding the other ratios fixed at unity. We find that the coexistence rescue is strongest under *k*_self_ *>* 1 (Fig. 4E) i.e., when the species 2 inflicts more non-consumptive interference on itself relative to species 1, sublinearity opens coexistence at niche overlaps where generalised Lotka-Volterra predicts exclusion. Analogously, species coexistence occurs under decreased interference by the dominant species (Fig. 4D) relative to species 1 interfering the dominant species 2.

### 3.4 Eco-evolutionary trajectories confirm coexistence at extreme niche overlap under sublinearity

To synthesise further the analytical and simulation results, we project the eco-evolutionary trajectories of two species as into MCT space (Fig. 5). With very similar mean trait values *µ*_1_ = 0.21; *µ*_2_ = 0.2, the eco-evolutionary trajectory of niche overlap and fitness ratio (*ρ* → 1, *κ*_1_/*κ*_2_ → 1) tends towards 1. At this point, species need to differentiate in order to coexist. We find that at low trait variance *σ*_1_ = 0.01 for both species (see table S1 for details), and for both Lotka-Volterra and sublinear trajectories (*γ* = 1; *γ* = 0.5), dynamically evolved towards the coexistence parameter space (designated by grey space) and evolutionary trait divergence results in a higher niche difference, ending inside the coexistence region (Fig. 5). However, we find the result is surprising for high trait variance. With initial parameter values (*µ*_1_ = 0.21; *µ*_2_ = 0.2), both species start outside the coexistence region initially since they have nearly identical trait means. Over eco-evolutionary time we observed that evolutionary differentiation fails under generalised Lotka-Volterra model: evolutionary endpoint lies outside the coexistence region and one species goes extinct (the trajectory exits toward *κ*_1_/*κ*_2_ → 0). However, under sublinearity due to species interference (*γ* = 0.5; *γ* = 1), the same evolutionary endpoint lies inside the widened coexistence region, and both species coexist ((Fig. 5, insets). This is the visual signature of the species coexistence rescue. That is, an eco-evolutionary community sitting at a MCT space where Lotka-Volterra model predicts exclusion but sublinear density dependence permits coexistence. In the sublinear model using the same parameters (Fig. 5B), both species coexist and converge on nearly identical trait distributions. This demonstrates that species coexistence can occur without evolutionary trait divergence, shifting the system towards the neutrality continuum (26).

**Figure 5.**
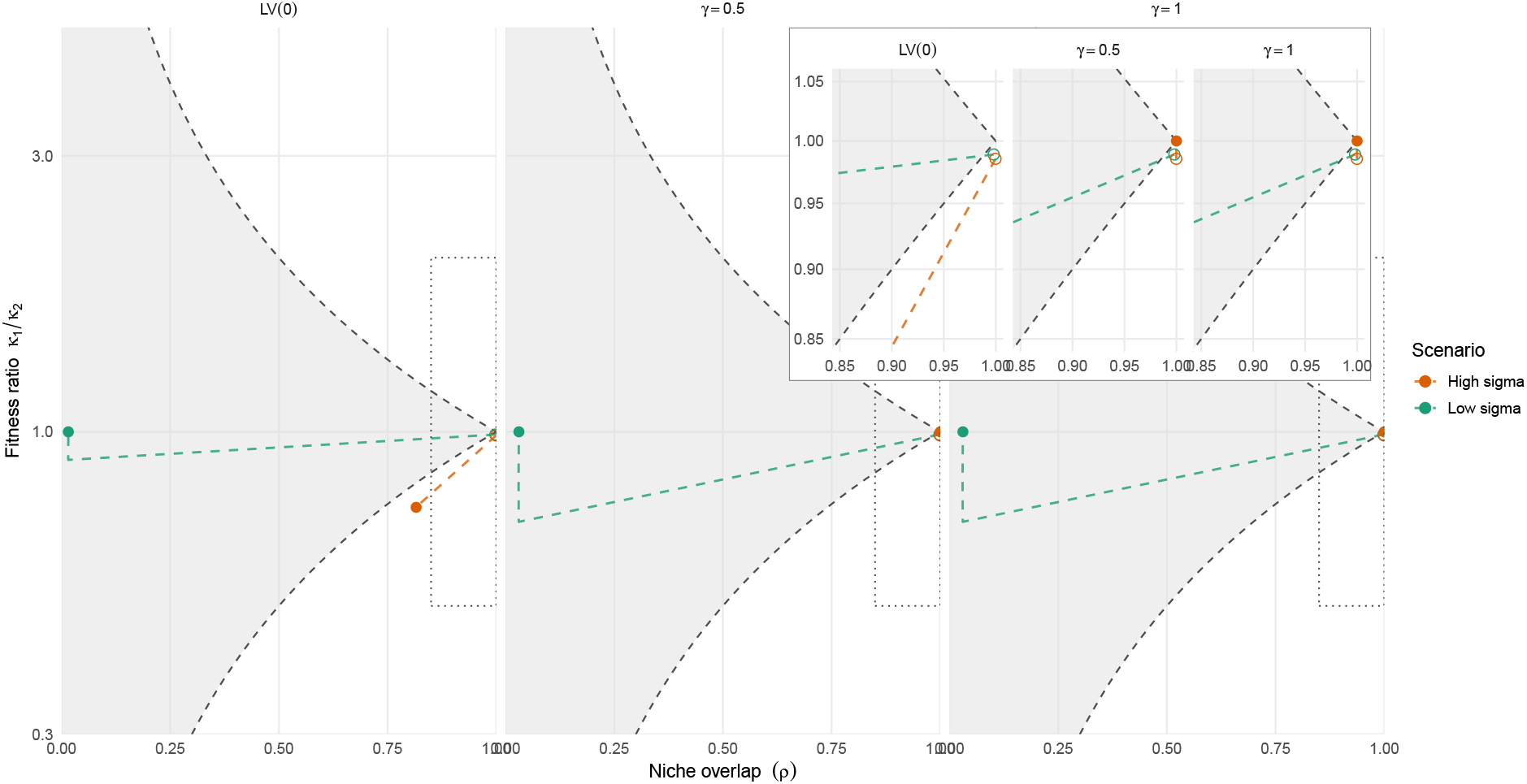
Coexistence at eco-evolutionary equilibria represented by the grey space designated by fitness ration *κ*_1_/*κ*_2_ and niche overlap *ρ* under interfence. Trajectories show the joint eco-evolutionary dynamics of niche overlap *ρ* and fitness ratios for two competiting species projected on the MCT coexistence plane (equations 23-24). The grey area designates coexistence region domain, and trajectories leaving this region indicate competitive exclusion. Panels/columns contrast generalised trait-based LV dynamics *c*_*i*_ = 0, and with interference *γ* = 0.5, *c*_*i*_ = 1 (nearest neighbour mechanism) and with *γ* = 1, *c*_*i*_ = 1 (well-mixed interference), for low *σ*_*i*_ = 0.01 and high *σ*_*i*_ = 0.10 trait variance. Open and filled circles mark the start *t* = 0, and end of eco-evolutionary simulations *t* = 10^5^ for each trajectory. Species were initialised with nearly identical trait means *µ*_1_ = 0.21, *µ*_2_ = 0.20, so that all trajectories start with high niche overlap. Under low trait standard deviation (green color dashed line), evolutionary trait divergence drives the mean traits of the two species apart reducing niche overlap and the two species move towards the grey space of coexistence region with minimal niche overlap in all three models. Under high trait variance, and under generalised LV dynamics, trait divergence fails and one competitive exclusion occurs (orange colour dashed line). Under interference, *γ* = 1, 0.5, both species persist but at high niche overlap (insets, orange colors). Parameters used *w* = 0.1, *θ* = 0.75, *m*_0_ = 0.25, 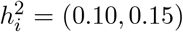, *c*_*i*_ = 1, *η*_*ii*_ = 1, *η*_*ij*_ = 0.01

### 3.5 Eco-evolutionary emergence of species coexistence in large competitive communities despite high intraspecific variation

We extend our MCT results of two species to larger competitive communities comprising of *S* = 30, 40, 50 species along a one-dimensional trait axis whose mean trait values *µ*_*i*_’s were randomly sampled from auniform distribution from − 0.7, to 0.7, and trait variances (high or low) i.e., *σ*_*i*_’s are also randomly sampled from a uniform distribution (see table S1), and we do this for three different models: LV, and the sublinear models (*γ* = 0.5, 1), and for varying levels of inter-intraspecific interference ratio, *ψ*. We found that sub-linear density dependence strongly promotes species coexistence at large competitive communities at eco-evolutionary equilibrium even at high species trait variation (Fig. 6, Fig. S2), and also leads to trait clustering (Fig. S3). We also found that as intraspecific interference coefficients become stronger relative to interspecific interference coefficient given by rising *ψ* values, species coexistence increased while in LV species coexistence was disrupted as has been stated in various previous studies (Fig. 6) (38; 39).

**Figure 6.**
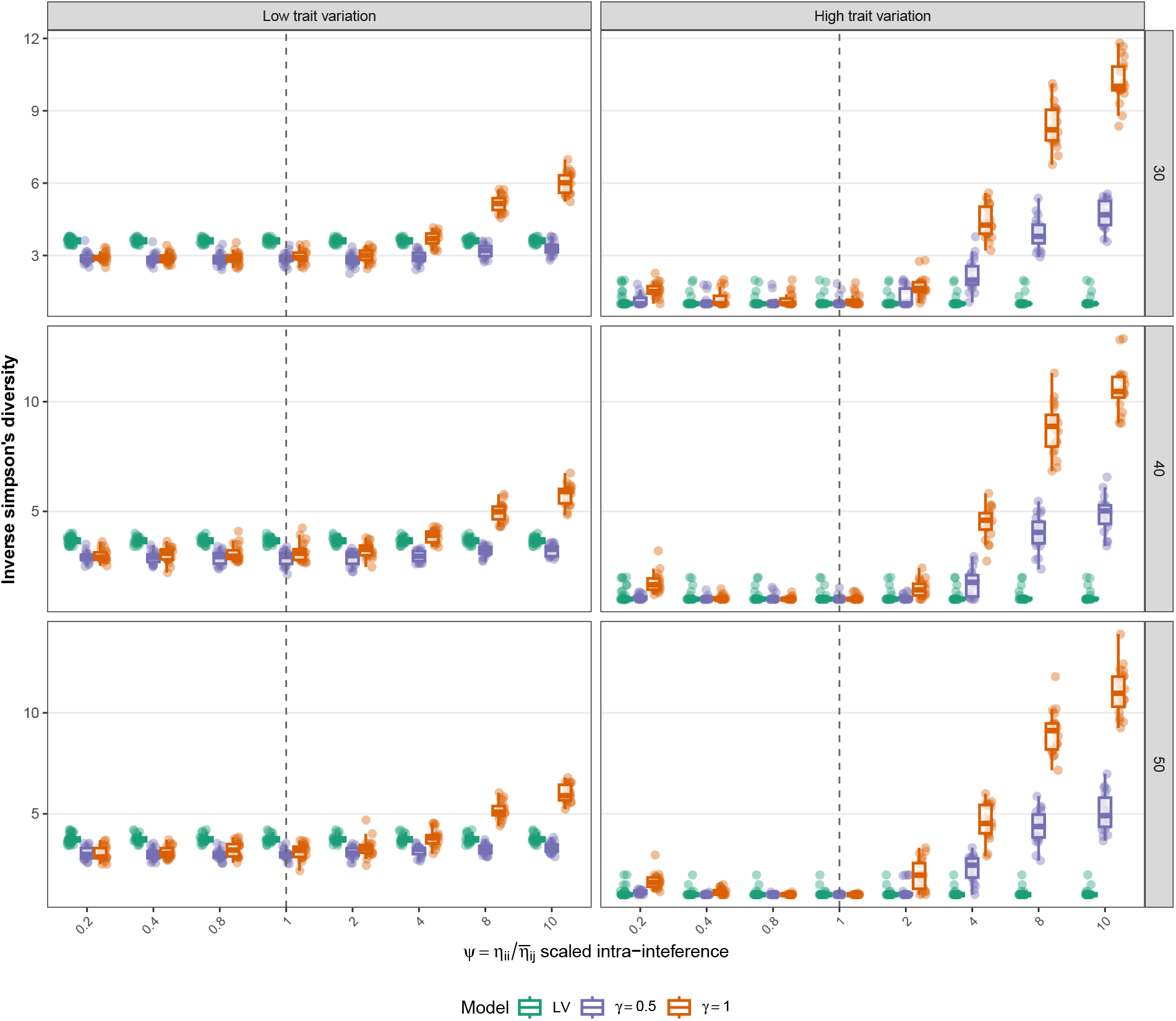
Species coexistence in large competitive communities in sublinear density regime. For initial *S* = 30, 40, 50 species competitive community (row panels), in eco-evolutionary equilibria species diversity (inverse Simpson’s index) was plotted against the ratio of intra-interference versus inter-interference factor, i.e., 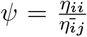, for sublinear models with *γ* = 0.5 (violet color boxplots), *γ* = 1 (orange color boxplots), and generalised Lotka-Volterra model (green boxplots). *γ* = 0.5 represents nearest-neigbour interference mechanism, and *γ* = 1 represents well-mixed interference mechanism. For generalised trait-based LV model, high trait variance (right figure column) leads to lower amount of species coexisting across different initial species community in contrast to low trait variation and sublinear models. Boxplots consists of 40 independent replicate simulations of our eco-evolutionary model (equation 21, 22). Parameters include from table S1.

## 4 Discussion

Recent studies have demonstrated a decelerating density-dependence pattern that has been supposedly observed across different taxonomic groups (15; 46). This density-dependence pattern has been subjected to debate as such a pattern can be observed in observational timeseries due to statistical issues (24). Nonetheless, such non-linear density dependence could have profound impacts on ecological stability, as increasing species diversity actually increases stability under sublinear density-dependence, which goes against the complexity-stability paradox (15; 47). This phenomenological observation of sublinear density dependence and its mechanistic emergence remains poorly understood and highly controversial. Classic and standard ecological models struggle to accurately represent sublinear density dependence (20; 23). When formulated phenomenologically, these models suffer from a critical mathematical flaw: per-capita growth rates approach infinity at near-zero population densities, falsely implying that species would not go extinct (20; 23). If this flaw is artificially corrected, these models simply revert to standard competitive exclusion, where species richness undermines ecological stability (23). Consequently, the mechanistic origins of sublinear density dependence remain unclear. Resolving these controversies requires a mathematically rigorous framework capable of definitively demonstrating whether such non-linear dynamics can naturally emerge. Here, we presented a novel framework where we take a geometric approach to consumer-resource theory, and show that sublinearity can mechanistically emerge from simple equations based on geometric constraints. We further show using an individual based model, that was developed from first principles, that sublinear density dependence, or superlinearity-like behaviour can readily emerge when consumers exhibit non-consumptive interference that are characteristic of certain spatially aggregated behaviour to spatially segregated behaviour. Adding to this, we further qualitatively validate our findings using empirical data of consumer-resource experiments and show that sub-linearity can leave signatures in such experiments. Finally, we show that such sublinear density dependence emerging from non-consumptive interference systematically changes the coexistence regime and reverses the commonly destabilizing effect of intraspecific trait variation.

Embedding an interference-modified attack rate within logistic consumer-resource equations generates a previously unrecognised asymmetry. This asymmetry is that the consumer intrinsic growth rate is attenuated by a non-consumptive factor, *Q*(*N*), whereas resource-mediated density-dependence is attenuated by *Q*(*N*)^2^. This attenuation arises due to consumers engaging in non-consumptive interactions which has consequences both in demographic rates as well as in competition. The shape of this *Q*(*N*), however, emerges when we take a geometric approach of consumers confined in an arena of a certain area. This is very generalas species in nature usually reside in a certain area where they forage. From that perspective, if consumers were confined in a certain habitat, non-consumptive interactions, and interactions arising due to resource competition will have disproportionate role in shaping density-dependence. Indeed, taking such a geometric view of non-consumptive interactions readily leads to mechanistic derivation of *Q*(*N*) which turns out to be non-linear in nature. Extending this perspective further, consumers in a habitat of a certain area will spatially move around for resources. This will directly generate different spatial behaviours, which could have consequences on how individuals interfere with each other. For instance, certain individuals might prefer to forage in their local neighbourhood vicinity. This would lead to non-consumptive interference at the immediate local scale. This is akin to the scenario where interference is determined by a nearest-neighbour mechanism, i.e., individuals interfere non-consumptively more so in their neighbourhood than afar. This mechanism produces an interference rate that scales as *N* ^1/2^. Such scaling may also arise when consumers exhibit spatially restricted movement (48), territorial behaviour (49), local aggregation (50; 51; 52) and neighbour interactions (29; 53), crowding behaviour (54; 55), or directed responses towards nearby conspecifics, causing interference to be governed primarily by nearest-neighbour distances. Consistent with this prediction, our individual-based model simulations show that when consumers have an affinity towards their immediate neighbours leading to spatial aggregation, the estimated interference exponent is approximately around *γ* ≈ 1/2 (Fig. 2, Fig. S4), in agreement with the analytical model derived in Box 1 and Box 2, but with some variability (Fig. S4).

Contrary to this, consumers can also exhibit free movement behaviour in a habitat. In such a scenario, when individual consumers move freely in a well-mixed scenario, rate of interference scales roughly linearly to the density in the area (40; 41; 56; 57). As a result, we get a different *Q*(*N*) where the interference exponent comes out analytically to be *γ* = 1. We further show that such a interference exponent emerges from our individual-based simulation results, where we ensure that consumers randomly move around in the arena, and that interference can occur through random movement. This interference which we call the “well-mixed interference mechanism” is of the form of classic Beddington-DeAngelis interference curve (40; 41). Beddington-DeAngelis time-budget interpretation of consumer interference (our well-mixed interference regime,*γ* = 1) is based on predator time among searching, handling and interference, and their derivation does not emphasise on the demographic asymmetry that can arise, which we do here. To be also noted that our geometric approach of consumer interference was not to derive a new functional response, but to show that that alternative spatial encounter mechanisms generate different density-scaling exponents within a regularized interference framework. In addition, we show further that these exponents transform resource-mediated demographic regulation and thereby have significant implications on species coexistence.

Since, our interference mechanisms are general, and scales as *N*^*γ*^, our framework can also produce interference that leads to superlinear-like density dependence. This can possibly arise when *γ >* 1, and when consumers exhibit some kind of non-aggregate or avoiding-interference behaviour. Indeed, when consumers avoided conspecifics quantified by *ξ <* 0 in our individual-based model, density dependence remained weak at low population densities because individuals could successfully evade interference. However, as density increased, due to the finite area, progressively fewer escape routes were available. Consequently, this led to avoidance behaviour to fail and interference to increase subsequently. This threshold-like transition generated superlinear interference scaling (*γ >* 1), together with a steep reduction in realised feeding time and per-capita growth at high density (Fig. 2D). Recent microbial experiments illustrate that the inferred shape of density dependence depends on both the experimental regime and the density range examined (19; 21; 22; 25). Our framework identifies non-consumptive conspecific interference as one possible mechanism contributing to such non-resource regulation. It also, in addition, emphasises that the interference exponent describes the scaling of loss of feeding-time and thus is not directly equivalent to the curvature exponent demonstrated in those studies. Nonetheless, we instead argue that the debate between the presence of sublinear, linear, or superlinear density dependence may arise from treating population regulation as a single phenomenological relationship. Our framework does not do this. Instead it actively tries to separate the spatial scaling of conspecific encounters/interference from its demographic consequences, and then further shows that behavioural mechanistic interference, resource competition, and different resource-supply regimes can generate different, and sometimes sequential, regions of density-dependent curvature. Empirically, using consumer-prey experimental timeseries we were able to qualitatively show that interference is prevalent in such systems. Specifically, we tried to fit models with different interference exponent, *γ* = 0, 1, 1/2, and qualitatively evaluate which model with interference exponent fits best on the experimental data. Across empirical time-series analysed, we found moderate support that interference exists (60 % of data) and can possibly lie between *γ* = 1, 1/2 (Fig. 3). Our analysis of empirical results underscore the importance of interference mediated density-dependence that can be prevalent in such systems and thus potentially scale up to impact biodiversity. Indeed, previous studies have shown that non-linear density-dependence can be widespread (50; 51; 52) and have impact on ecosystem functions (50).

Embedding such a spatial and geometric mechanism of interference in competition theory reveals that interference acts on the fitness axis of modern coexistence theory while leaving the niche axis untouched. In our models with interference, niche overlap depends only on the competition coefficients and thus remained unchanged (32). What in addition we found was that the intrinsic growth rates were replaced by resident-species-conditional effective rates, so that the fitness ratio takes different values depending on which species was at equilibrium. This dependence on the resident species had a direct consequence: the coexistence region retains of finite width as niche overlap approaches one. However,under Lotka–Volterra model, it collapsed to a point and any asymmetry in demographic rates could potentially exclude one competitor (Fig. 4). Our results thus demonstrate that interference permits coexistence in precisely the regime where classical Lotka-Volterra stabilisation was not available. In addition, the direction of such an effect was governed by a simple condition — whether a rare species escapes more interference than the resident species inflicts interference upon itself. Our eco-evolutionary dynamical results further show when this matters (Fig. 5-6). Under low intraspecific trait variance, trait divergence generates niche differences and the two models converge towards the same result (32). However, under high trait variance our results demonstrated something different. We found that trait differentiation does not occur, and interference became the sole available path to coexistence. This was a very surprising and interesting result. Potentially, this makes intraspecific trait variation a potential switch between stabilising mechanisms rather than simply a modifier of competition strength. This has strong implications since it suggests that systems in which species appear ecologically similar need not be closer to competitive exclusion. Our results have the potential to explain why intraspecific variation is pervasive, and that it can reverse the classical result of trait variation disrupting species coexistence (37; 38; 39). When we extended this to multispecies communities, we observed similar results, that intraspecific trait variation can promote species coexistence in large multispecies competitive communities, contingent on the structure of the interference matrix relative to the competition matrix (Fig. 6). Our study suggests that many species can coexist when intra-interference is substantially stronger than inter-interference, indicating that non-consumptive interactions are not merely a correction to resource competition but a distinct axis along which diverse communities may be assembled.

The current debate on density-dependence indicates the presence of a dichotomy between resource-driven superlinearity and non-resource driven sublinearity (15; 19; 20; 21; 22; 23). Our framework from a spatial geometrical-interference provides a mechanistic bridge. We show that our geometric interference approach alters resource acquisition, produces regularised sublinear demographic feedback, and creases alternative axis of competition that can stabilise coexistence even when resource niches strongly overlap. Our framework shows that sublinear or superlinear density dependence are not necessarily competing descriptions or hypotheses. Instead, they can emerge under different mechanisms, and at different densities and under different resource regimes. Non-consumptive interference as shown in our study, can be one possible source of resource-independent inhibition, although it may not be necessarily the only one. Our study thus raises many further questions. Specifically, it raises the question of whether *γ* is a fixed characteristic of a population or whether if can change depending on the density gradient, behaviour, or environmental constraints. Habitat configuration, confinement, conspecific attraction, interaction distance, resource availability etc., can determine whether interference approaches the nearest-neighbour mechanism or well-mixed mechanism. These predictions can motivate experiments that directly quantify interference rates across densities to elucidate where the interference exponent changes, which has consequences on emergence of a density-dependent form that further can scale and impact biodiversity.

## 5 Acknowledgements

The authors thank György Barabás, and Meike Wittmann for their comments on the previous versions of this framework. We used Anthropic’s Claude Opus models for finding empirical data suitable to fit our framework. The authors declare no conflict of interests.

## 6 Data Availability

The data and code will be uploaded to Zenodo. R scripts for simulations are available in the repository https://github.com/GauravKBaruah/geometric-interactions-sublinear.

## References

[1] Clark, J. S. Individuals and the Variation Needed for High Species Diversity in Forest Trees. Science 327, 1129–1132 (2010). URL http://www.sciencemag.org/content/327/5969/1129.abstract.

[2] Singh, P. & Baruah, G. Higher order interactions and species coexistence. Theoretical Ecology (2020). URL 10.1007/s12080-020-00481-8.

[3] Barabás, G., Michalska-Smith, M. J. & Allesina, S. Self-regulation and the stability of large ecological networks. Nature Ecology & Evolution 1, 1870–1875 (2017). URL https://www.nature.com/articles/s41559-017-0357-6.

[4] Saavedra, S. et al. A structural approach for understanding multispecies coexistence. Ecological Monographs 87, 470–486 (2017). URL http://doi.wiley.com/10.1002/ecm.1263.

[5] Gravel, D., Guichard, F. & Hochberg, M. E. Species coexistence in a variable world. Ecology Letters 14, 828–839 (2011).

[6] Chesson, P. Mechanisms of Maintenance of Species Diversity. Annual Review of Ecology and Systematics 31, 343–366 (2000). URL http://www.annualreviews.org/doi/10.1146/annurev.ecolsys.31.1.343.

[7] Barabás, G., Meszéna, G. & Ostling, A. Community robustness and limiting similarity in periodic environments. Theoretical Ecology 5, 265–282 (2012). URL http://link.springer.com/10.1007/s12080-011-0127-z.

[8] Kremer, C. T. & Klausmeier, C. A. Coexistence in a variable environment: Eco-evolutionary perspectives. Journal of Theoretical Biology 339, 14–25 (2013).

[9] Suweis, S., Grilli, J., Banavar, J. R., Allesina, S. & Maritan, A. Effect of localization on the stability of mutualistic ecological networks. Nature Communications 6, 10179 (2015). URL https://www.nature.com/articles/ncomms10179.

[10] Rohr, R. P., Saavedra, S. & Bascompte, J. On the structural stability of mutualistic systems. Science 345 (2014). URL https://science.sciencemag.org/content/345/6195/1253497.

[11] Baruah, G. The impact of individual variation on abrupt collapses in mutualistic networks. Ecology Letters 25, 26–37 (2022). URL https://onlinelibrary.wiley.com/doi/abs/10.1111/ele.13895. _eprint: https://onlinelibrary.wiley.com/doi/pdf/10.1111/ele.13895.

[12] Stouffer, D. B. & Bascompte, J. Compartmentalization increases food-web persistence. Proceedings of the National Academy of Sciences 108, 3648–3652 (2011).

[13] Baruah, G. & Wittmann, M. Reviving collapsed plant–pollinator networks from a single species. PLOS Biology 22, e3002826 (2024). URL https://journals.plos.org/plosbiology/article?id=10.1371/journal.pbio.3002826. Publisher: Public Library of Science.

[14] Sibly, R. M. et al. Fundamental insights into ontogenetic growth from theory and fish. Proceedings of the National Academy of Sciences 112, 13934–13939 (2015). URL https://www.pnas.org/content/112/45/13934.

[15] Hatton, I. A., Mazzarisi, O., Altieri, A. & Smerlak, M. Diversity begets stability: Sublinear growth and competitive coexistence across ecosystems. Science 383, eadg8488 (2024).

[16] Allesina, S. & Tang, S. Stability criteria for complex ecosystems. Nature 483, 205–208 (2012). URL https://www.nature.com/articles/nature10832.

[17] Abrams, P. A. Determining the functional form of density dependence: deductive approaches for consumer-resource systems having a single resource. The American Naturalist 174, 321–330 (2009).

[18] Smith, F. E. Population dynamics in daphnia magna and a new model for population growth. Ecology 44, 651–663 (1963).

[19] Letten, A. D. Making sense of (sublinear) density dependence. Trends in Ecology & Evolution 40, 622–625 (2025).

[20] Fronhofer, E. A., Govaert, L., O’Connor, M. I., Schreiber, S. J. & Altermatt, F. The shape of density dependence and the relationship between population growth, intraspecific competition and equilibrium population density. Oikos 2024, e09824 (2024).

[21] Mazzarisi, O. et al. Universal sublinear population growth density dependence unrelated to resource limitation. bioRxiv 2025–12 (2025).

[22] Orr, J. A. et al. Growth-density inversion in escherichia coli reveals superlinear, not sublinear, and density dependence. PLoS biology 24, e3003898 (2026).

[23] Aguadé-Gorgorió, G., Lajaaiti, I.Arnoldi, J.-F. & Kéfi, S. Unpacking sublinear growth: diversity, stability and coexistence. Oikos 2025, e10980 (2025).

[24] Doncaster, C. P. Non-linear density dependence in time series is not evidence of non-logistic growth. Theoretical Population Biology 73, 483–489 (2008).

[25] Letten, A. D. et al. Shifting resource limitation explains multiphasic patterns of density dependence. bioRxiv 2026–07 (2026).

[26] Hubbell, S. P. Neutral Theory and the Evolution of Ecological Equivalence. Ecology 87, 1387–1398 (2006). URL 10.1890%2F0012-9658%282006%2987%5B1387%3ANTATEO%5D2.0.CO%3B2.

[27] Abrams, P. A. Arguments in Favor of Higher Order Interactions. The American Naturalist 121, 887–891 (1983). URL https://www.journals.uchicago.edu/doi/10.1086/284111.

[28] Grilli, J., Barabás, G., Michalska-Smith, M. J. & Allesina, S. Higher-order interactions stabilize dynamics in competitive network models. Nature 548, 210–210 (2017). URL http://www.nature.com/doifinder/10.1038/nature23273.

[29] Wiegand, T. et al. Latitudinal scaling of aggregation with abundance and coexistence in forests. Nature 640, 967–973 (2025).

[30] Chesson, P. Updates on mechanisms of maintenance of species diversity. Journal of Ecology 106, 1773–1794 (2018). URL http://doi.wiley.com/10.1111/1365-2745.13035.

[31] Letten, A. D.Ke, P.-J. & Fukami, T. Linking modern coexistence theory and contemporary niche theory. Ecological Monographs 87, 161–177 (2017). URL http://doi.wiley.com/10.1002/ecm.1242.

[32] Pastore, A. I., Barabás, G., Bimler, M. D., Mayfield, M. M. & Miller, T. E. The evolution of niche overlap and competitive differences. Nature Ecology & Evolution 5, 330–337 (2021). URL https://www.nature.com/articles/s41559-020-01383-y. Number: 3 Publisher: Nature Publishing Group.

[33] Siefert, A. et al. A global meta-analysis of the relative extent of intraspecific trait variation in plant communities. Ecology Letters 18, 1406–1419 (2015). URL http://doi.wiley.com/10.1111/ele.12508.

[34] Siefert, A. Incorporating intraspecific variation in tests of trait-based community assembly. Oecologia 170, 767–775 (2012). URL http://link.springer.com/10.1007/s00442-012-2351-7.

[35] Clark, J. S. et al. High-dimensional coexistence based on individual variation: A synthesis of evidence. Ecological Monographs 80, 569–608 (2010).

[36] Violle, C. et al. The return of the variance: intraspecific variability in community ecology. Trends in Ecology & Evolution 27, 244–252 (2012). URL https://www.sciencedirect.com/science/article/pii/S0169534711003375.

[37] Hart, S. P., Schreiber, S. J., Levine, J. M. & Coulson, T. How variation between individuals affects species coexistence. Ecology Letters 19, 825–838 (2016).

[38] Barabas, G. & D’Andrea, R. The effect of intraspecific variation and heritability on community pattern and robustness. Ecology Letters 19, 977–986 (2016).

[39] Baruah, G., Barabás, G. & John, R. When Do Trait-Based Higher Order Interactions and Individual Variation Promote Robust Species Coexistence? Ecology and Evolution 15, e71336 (2025). URL https://onlinelibrary.wiley.com/doi/abs/10.1002/ece3.71336. _eprint: https://onlinelibrary.wiley.com/doi/pdf/10.1002/ece3.71336.

[40] Beddington, J. R. Mutual interference between parasites or predators and its effect on searching efficiency. The Journal of Animal Ecology 331–340 (1975).

[41] DeAngelis, D. L., Goldstein, R. A. & O’Neill, R. V. A model for tropic interaction. Ecology 56, 881–892 (1975).

[42] Skalski, G. T. & Gilliam, J. F. Functional responses with predator interference: viable alternatives to the holling type ii model. Ecology 82, 3083–3092 (2001).

[43] Uiterwaal, S. F., Lagerstrom, I. T., Lyon, S. R. & DeLong, J. P. Forage database: A compilation of functional responses for consumers and parasitoids (2022).

[44] Rosenzweig, M. L. & MacArthur, R. H. Graphical representation and stability conditions of predator-prey interactions. The American Naturalist 97, 209–223 (1963).

[45] O’Dwyer, J. P. Whence lotka-volterra? conservation laws and integrable systems in ecology. Theoretical Ecology 11, 441–452 (2018).

[46] Sibly, R. M., Barker, D., Denham, M. C., Hone, J. & Pagel, M. On the regulation of populations of mammals, birds, fish, and insects. Science 309, 607–610 (2005).

[47] May, R. M. Qualitative Stability in Model Ecosystems. Ecology 54, 638–641 (1973). URL http://doi.wiley.com/10.2307/1935352.

[48] Inchausti, P. & Ballesteros, S. Intuition, functional responses and the formulation of predator–prey models when there is a large disparity in the spatial domains of the interacting species. Journal of animal ecology 77, 891–897 (2008).

[49] Grant, J. W., Weir, L. K. & Steingrímsson, S. Ó. Territory size decreases minimally with increasing food abundance in stream salmonids: Implications for population regulation. Journal of Animal Ecology 86, 1308–1316 (2017).

[50] Little, C. J., Fronhofer, E. A. & Altermatt, F. Nonlinear effects of intraspecific competition alter landscape-wide scaling up of ecosystem function. The American Naturalist 195, 432–444 (2020).

[51] Kullmann, H., Thünken, T., Baldauf, S. A., Bakker, T. C. & Frommen, J. G. Fish odour triggers conspecific attraction behaviour in an aquatic invertebrate. Biology Letters 4, 458 (2008).

[52] Van Gils, J. A. & Piersma, T. Digestively constrained predators evade the cost of interference competition. Journal of Animal Ecology 73, 386–398 (2004).

[53] Baruah, G., Molau, U., Jägerbrand, A. K. & Alatalo, J. M. Impacts of seven years of experimental warming and nutrient addition on neighbourhood species interactions and community structure in two contrasting alpine plant communities. Ecological Complexity 33, 31–40 (2018). URL https://www.sciencedirect.com/science/article/pii/S1476945X17301101.

[54] Venkitachalam, S., Sajith, V. & Joshi, A. More than just density: the role of egg number, food volume and container dimensions in mediating larval competition in drosophila melanogaster. bioRxiv 2023–07 (2023).

[55] Venkitachalam, S. & Joshi, A. An individual-based simulation framework exploring the ecology and mechanistic underpinnings of larval crowding in laboratory populations of drosophila. Journal of Theoretical Biology 112378 (2026).

[56] Van Der Meer, J. & Ens, B. J. Models of interference and their consequences for the spatial distribution of ideal and free predators. Journal of Animal Ecology 846–858 (1997).

[57] Hassell, M. & Varley, G. New inductive population model for insect parasites and its bearing on biological control. Nature 223, 1133–1137 (1969).

