## appendix for "Geometric scaling of non-consumptive interactions generates sublinear density dependence and reshapes coexistence"

#### 1 Single species equilibrium with interference:

$$\frac{dN_i}{dt} = N_i\big[b_iQ_i - m_0 - Q_i\big(Q_i\alpha_{ii}N_i + Q_j\alpha_{ij}N_j\big)\big], \quad i,j \in \{1,2\}, \tag{S1}$$

with the species-specific interference factor following from BOX 1,

$$Q_i = \frac{1}{1 + c_i\big[\eta_{i1}N_1 + \eta_{i2}N_2\big]^{\gamma}} \tag{S2}$$

Where $\gamma$ gives the interference scaling factor that emerges due to consumer movement behaviour (Box 1 and Box 2). In single species only the intraspecific term of Eq. S1 survives, so the equilibrium density $K_i$ satisfies

$$b_iQ_i^* - m_0 = (Q_i^*)^2\,\alpha_{ii}K_i, \qquad Q_i^* = \frac{1}{1 + c_i(\eta_{ii}K_i)^{\gamma}}. \tag{S3}$$

Substituting $d_i = c_i(\eta_{ii}K_i)^{\gamma}$, so that $Q_i^* = (1 + d_i)^{-1}$ and $K_i = \eta_{ii}^{-1}(d_i/c_i)^{1/\gamma}$, and multiplying through by $(1 + d_i)^2$ gives:

$$b_i(1 + d_i) - m_0(1 + d_i)^2 = \frac{\alpha_{ii}}{\eta_{ii}}\left(\frac{d_i}{c_i}\right)^{1/\gamma}. \tag{S4}$$

For $\gamma = 1$ this is a quadratic in $u_i$ with the unique positive root

$$d_i = \frac{B_i + \sqrt{B_i^2 + 4m_0(b_i - m_0)}}{2m_0}, \qquad B_i = b_i - 2m_0 - \frac{\alpha_{ii}}{c_i\eta_{ii}}, \tag{S5}$$

and for general $\gamma$ Eq. S4 is solved numerically. A positive solution exists whenever $b_i > m_0$, which is the same persistence condition as under Lotka-Volterra dynamics. Now, note the equilibrium identity is written as from eq. S3:

$$\alpha_{ii}K_i = \frac{b_iQ_i^* - m_0}{(Q_i^*)^2}. \tag{S6}$$

##### 1.1 Invasion analysis

During invasion analysis, we let species $i$ be rare while species $j$ rests at $K_j$. The invader experiences interference generated solely by the resident species, and the resident species depletion of the shared resource is attenuated by its own interference factor. Hence we can write it as:

$$\bar{r}_i = b_iQ_i^{(j)} - m_0 - Q_i^{(j)}Q_j^*\,\alpha_{ij}K_j, \qquad Q_i^{(j)} = \frac{1}{1 + c_i(\eta_{ij}K_j)^{\gamma}}. \tag{S7}$$

Dividing $\bar{r}_i > 0$ by $Q_i^{(j)} > 0$, and substituting $\alpha_{ij}K_j = (\alpha_{ij}/\alpha_{jj})\,\alpha_{jj}K_j$ and then applying Eq. S6 to the resident species, we obtain:

$$\bar{r}_i > 0 \quad\Longleftrightarrow\quad \frac{\tilde{R}_i^{(j)}}{\tilde{R}_j^{(j)}} > \frac{\alpha_{ij}}{\alpha_{jj}}, \tag{S8}$$

where the resident species conditional effective rate is given as:

$$\tilde{R}_i^{(j)} \equiv \frac{b_iQ_i^{(j)} - m_0}{Q_i^{(j)}} = b_i - \frac{m_0}{Q_i^{(j)}} = \underbrace{\big(b_i - m_0\big)}_{r_i} - \underbrace{m_0\,c_i\big(\eta_{ij}K_j\big)^{\gamma}}_{\text{interference penalty}}. \tag{S9}$$

We find that the Equation S8 is structurally identical to the Lotka-Volterra invasion criterion $r_i/r_j > \alpha_{ij}/\alpha_{jj}$, with $\tilde{R}$ replacing $r$. Equation S9 shows that non-consumptive interference acts as a penalty on the intrinsic rate, proportional to the density-independent mortality $m_0$ and scaling as the $\gamma$-power of the crowding the invader receives from the resident.

##### 1.2 Resource dynamics and the quasi-equilibrium resource and derivation of two species equations:

We consider two consumer species with densities $N_1$ and $N_2$ feeding on a single resource $R$. The resource grows at intrinsic rate $p$, self-regulates/limits at rate $\alpha$, and is depleted by both consumers at utilisation rate $u$,

$$\frac{dR}{dt} = pR - \alpha R^2 - u\,(N_1 + N_2)\,R \tag{S10}$$

We follow the classical assumption that the resource turnover is rapid relative to consumer population dynamics[1,2], we set $dR/dt = 0$ and divide through by $R > 0$ to obtain the quasi-equilibrium resource level

$$R^* = \frac{p}{\alpha} - \frac{u}{\alpha}\,(N_1 + N_2). \tag{S11}$$

The resource thus declines linearly in the total consumer density: each additional consumer, of either species, draws the shared resource further down of course. We allow the utilisation rate to differ among consumers, $u \to u_j$ for species $j$, and generalising to $S$ species gives

$$R^* = \frac{p}{\alpha} - \sum_{j=1}^{S} \frac{u_j}{\alpha}\,N_j = R_0 - \sum_{j=1}^{S} u'_j\,N_j, \tag{S12}$$

where $R_0 = p/\alpha$ is the resource saturation level in the absence of consumption and $u'_j = u_j/\alpha$ are the rescaled utilisation rates. Equation S12 is the multispecies analogue of Eq. S11: the resource is its consumer-free level minus the summed consumption contributed by every consumer species present.

##### 1.3 Consumer dynamics without interference

The per-capita growth rate of consumer $i$ is proportional to its resource intake, converted into offspring with efficiency $\epsilon_i$, and a density-independent mortality $m_0$, so that it gives:

$$\frac{dN_i}{dt} = N_i\,r_i = N_i\big(\epsilon_i u_i R^* - m_0\big). \tag{S13}$$

Substituting Eq. S12 into Eq. S13 and expanding,

$$\frac{dN_i}{dt} = N_i\left(\epsilon_i u_i\Big(R_0 - \sum_{j=1}^{S} u'_j N_j\Big) - m_0\right) = \underbrace{\epsilon_i u_i R_0}_{b_i} N_i - \sum_{j=1}^{S} \underbrace{\epsilon_i u_i u'_j}_{\alpha_{ij}} N_j N_i - m_0 N_i, \tag{S14}$$

which is exactly the Lotka-Volterra competition form

$$\frac{dN_i}{dt} = N_i\left(b_i - m_0 - \sum_{j=1}^{S} \alpha_{ij} N_j\right), \tag{S15}$$

with intrinsic birth rate $b_i = \epsilon_i u_i R_0$ and competition coefficients $\alpha_{ij} = \epsilon_i u_i u'_j$.

The structure of $\alpha_{ij}$ is central. We set the conversion efficiency aside, abd then it is a product of two distinct utilisation rates: the focal species' uptake rate $u_i$ and the competitor species utilisation rate $u'_j$. Competition is thus not a single process but the composition of two i.e., how strongly $j$ utilises the resource, and how strongly $i$ depends on what is left. This factorisation is the single-resource counterpart of the trait-based competition kernel used in the main text. There, individuals are characterised by a phenotype $z$ and resources by a quality $y$, with $u(z, y)$ giving the rate at which phenotype $z$ utilises resource $y$; the competition kernel between phenotypes follows as the convolution (which we detail below in section 2),

$$\alpha(z, z') = \int u(z, y)\,u(y, z')\,\mathrm{d}y, \tag{S16}$$

which is again a product of the focal phenotype's utilisation and the competitor's utilisation, now resolved across the resource axis rather than collapsed onto a single resource. The two-rate structure is identical in both formulations, and it is this structure that dictates where the interference terms enter.

##### 1.4 Introducing non-consumptive interference

We now apply the same reasoning as in the single-species case (Box 1 and Box 2 in maint-ext). Interference among individuals reduces the fraction of time a consumer spends feeding. Species $i$'s realised utilisation rate is thus attenuated to $u_i Q_i$, where we get:

$$Q_i = \frac{1}{1 + c_i(\sum_j \eta_{ij} N_j)^\gamma} \in (0, 1] \tag{S17}$$

is the fraction of time species $i$ spends feeding, $c_i$ its interference sensitivity, and $\eta_{ij}$ the contribution of species $j$ to the crowding experienced by species $i$, and $\gamma$ is the interference mechanism scaling factor, 0.5 for nearest-neighbour and 1 for well-mixed interference (Box 1 and 2). Interference is non-consumptive: it does not itself remove resource from the system, but reduces the rate at which individuals can access what is there.

Importantly, we model this to be an attenuation that applies to each species in its own right. Species $j$'s depletion of the shared resource is reduced by $Q_j$, so Eq. S12 becomes which is given as:

$$R^* = R_0 - \sum_{j=1}^{S} u'_j\, Q_j\, N_j, \tag{S18}$$

while species $i$'s uptake of the remaining resource is separately reduced by $Q_i$, so Eq. S13 becomes

$$\frac{dN_i}{dt} = N_i\big(\epsilon_i u_i Q_i R^* - m_0\big). \tag{S19}$$

Substituting Eq. S18 into Eq. S19 and expanding as before,

$$\frac{dN_i}{dt} = N_i\left(\epsilon_i u_i Q_i R_0 - m_0 - \epsilon_i u_i Q_i \sum_{j=1}^{S} u'_j Q_j N_j\right) = N_i\left(b_i Q_i - m_0 - Q_i \sum_{j=1}^{S} Q_j\, \alpha_{ij} N_j\right). \tag{S20}$$

The two interference factors enter at different points and for different reasons. The factor $Q_i$ multiplying $b_i$ arises once, from the reduction in the focal species' own intake. The competition term acquires two factors, $Q_i$ from the focal species' reduced uptake and $Q_j$ from competitor $j$'s reduced depletion of the shared resource. Thus, one factor for each of the two utilisation rates whose product constitutes $\alpha_{ij}$ (Eq. S14). Setting $c_i = 0$ for all species gives $Q_i \equiv 1$ and recovers generalised Lotka-Volterra equations Eq. S15.

The same thing carries over to the trait-explicit formulation [(2)]. As each utilisation rate in the convolution of Eq. S16 is attenuated by the interference experienced by the corresponding phenotype, the trait-level competition term becomes and is given as:

$$Q_i(z, \mathbf{N}) \sum_{j=1}^{S} N_j \int Q_j(z', \mathbf{N})\, \alpha(z, z')\, p_j(z')\, \mathrm{d}z', \tag{S21}$$

with the competitor's interference factor appearing inside the integral over $z'$, since it is phenotype $z'$ whose depletion of the resource is reduced. Having written this we do not model that the same trait is responsible for resource competition and non-consumptive interference. While this is entirely possible, it is best left to understand how the structure of interference matrix (not dependent on any trait) can influence coexistence patterns in a trait-axis that influences resource competition. This is what we do in our study here. We however leave this same trait impacting non-consumptive interference and resource competition for future exploration.

##### 1.5 The two-species case

We write Eq. S20 out for $S = 2$ and then separate the intraspecific from the interspecific term, which gives:

$$\frac{dN_1}{dt} = N_1\big(b_1 Q_1 - m_0 - Q_1^2\, \alpha_{11} N_1 - Q_1 Q_2\, \alpha_{12} N_2\big), \tag{S22}$$

$$\frac{dN_2}{dt} = N_2\big(b_2 Q_2 - m_0 - Q_2^2\, \alpha_{22} N_2 - Q_2 Q_1\, \alpha_{21} N_1\big), \tag{S23}$$

with

$$Q_1 = \frac{1}{1 + c_1\big(\eta_{11} N_1 + \eta_{12} N_2\big)^\gamma}, \qquad Q_2 = \frac{1}{1 + c_2\big(\eta_{21} N_1 + \eta_{22} N_2\big)^\gamma}. \tag{S24}$$

The intraspecific terms carry $Q_i^2$ as the special case $j = i$ of the general rule: when a species competes with itself, both the uptake and the depletion factors refer to the same species and the product collapses to a square. The interspecific terms retain the distinct product $Q_1 Q_2$. In addition, $\gamma = 1, 0.5$ defines the interference mechanisms explained in figure 1 of main-text and BOX 1 and BOX 2. Since $Q_1$ and $Q_2$ respond to different crowding environments: species 1 feels $\eta_{11}N_1 + \eta_{12}N_2$ while species 2 gets $\eta_{21}N_1 + \eta_{22}N_2$ : the interspecific competition term is attenuated by a factor that is generally not equal to the attenuation of either species' intraspecific term. It is actually this asymmetry between $Q_i^2$ and $Q_iQ_j$ that generates the resident-dependent fitness ratios derived in the main text, and hence the departure of the coexistence region from its Lotka-Volterra counterpart.

#### 2 Trait-based generalisted Lotka-Volterra dynamics:

The species-level formulation does not capture the within-species phenotypic variation that mediates resource use. Next, we model phenotype-based McArthurs consumer resource eqaution that incorporates trait structure into by reconciling quantitative genetics framework into Lotka-Volerra equations. We assume individuals of consumers be characterised by a quantitative phenotype $z$ that determines how efficiently they use a resource of quality $y$, and is assumed to be normally distributed. IN such a case, the per-capita growth rate of phenotype $z$ in species $i$ is given as (see[2,3]:

$$r_i(z) = \int u(z, y)\, R(y)\, dy - m_0, \tag{S25}$$

following **(author?)**[1] and recent eco-evolutionary extensions[2,3]. The resource utilisation kernel $u(z, y)$ is Gaussian, $u(z, y) \propto \exp(-(z - y)^2/w^2)$, so that phenotype $z$ consumes most efficiently the resources matching its trait value. Under fast resource turnover, the resource concentration reaches quasi-equilibrium given as:

$$R(y) = R_0(y) - \sum_{j=1}^{2} N_j \int u(z', y)\, p_j(z')\, dz', \tag{S26}$$

where $p_j(z')$ is the phenotype distribution of species $j$, assumed Gaussian with mean $\mu_j$ and variance $\sigma_j^2$. Substituting Eq. S26 into Eq. S25 yields the Lotka-Volterra form given as:

$$r_i(z) = b_i(z) - \sum_{j=1}^{2} N_j \int \alpha(z, z')\, p_j(z')\, dz', \tag{S27}$$

where $\alpha(z, z') = \exp(-(z - z')^2/w^2)$ is the trait-based competition kernel between phenotypes[2,4]. To introduce sub-linearity at the trait level, we modify the utilisation kernel to account for phenotype-independent interference:

$$u_{\text{eff}}(z, y, \mathbf{N}) = Q_i(\mathbf{N})\, u(z, y), \tag{S28}$$

where the phenotype independependent interference factor is:

$$Q_i(z, \mathbf{N}) = \frac{1}{1 + c_i\left(\sum_{j=1}^{2} N_j \eta_{ij}\right)^{\gamma}}, \tag{S29}$$

With Eq. S29, the trait-based per-capita growth rate of phenotype $z$ in species $i$ becomes

$$r_i(z) = b_i(z)\, Q_i(\mathbf{N}) - m_0 - Q_i(\mathbf{N}) \sum_{j=1}^{2} Q_j(\mathbf{N}) N_j \int \alpha(z, z')\, p_j(z')\, dz'. \tag{S30}$$

Integrating Eq. S30 over the trait distribution $p_i(z)$ yields the species-level population dynamics given as:

$$\frac{dN_i}{dt} = N_i \int r_i(z) p_i(z) dz = N_i \left[ \overline{bQ_i} - m_0 - Q_i \sum_{j=1}^{S} Q_j \alpha_{ij} N_j \right], \tag{S31}$$

and evolutionary dynamics of the mean phenotype is given as[2]:

$$\frac{d\mu_i}{dt} = h_i^2 \int (z - \mu_i)\, r_i(z)\, p_i(z)\, dz = h_i^2 \left[ g_i - Q_i \sum_{j=1}^{S} Q_j \varphi_{ij} N_j \right], \tag{S32}$$

Where the competition coefficient is given as

$$\alpha_{ij} = \frac{w}{\sqrt{2\sigma_i^2 + 2\sigma_j^2 + w^2}} \exp\left( -\frac{(\mu_i - \mu_j)^2}{2\sigma_i^2 + 2\sigma_j^2 + w^2} \right), \tag{S33}$$

where $h_i^2$ is the heritability of the focal trait. Under the Gaussian trait assumption and the quadratic growth function, this reduces to with the trait selection coefficient due to competition as

$$\varphi_{ij} = \alpha_{ij} \frac{2\sigma_i^2(\mu_j - \mu_i)}{2\sigma_i^2 + 2\sigma_j^2 + w^2}. \tag{S34}$$

#### 3 Trait-based niche overlap and fitness differences undernon-consumptive interference

We derive the modern coexistence theory (MCT) decomposition for the trait-based model of the main text following from Pastore *et al*[(4)]. We show that the niche overlap $\rho$ is unaffected by interference, whereas the fitness ratio becomes resident-species-conditional, and we give explicit expressions for both in terms of the trait means $\mu_i$, trait variances $\sigma_i^2$, the competition kernel width $w$, and the interference matrix $\boldsymbol{\eta}$. This section is used to produce figure 5 i.e., the evolution of niche overlap and fitness difference under interference and under generalised Lotka-Volterra models.

We assume that individuals of species $i$ has a quantitative phenotype $z$ distributed as $p_i(z) \sim \mathcal{N}(\mu_i, \sigma_i^2)$. The per-capita growth rate of phenotype $z$ is

$$r_i(z) = b_i(z)\, Q_i(\mathbf{N}) - m_0 - Q_i(\mathbf{N}) \sum_{j=1}^{S} Q_j(\mathbf{N})\, N_j \int \alpha(z, z')\, p_j(z')\, \mathrm{d}z', \tag{S35}$$

with competition kernel $\alpha(z, z') = \exp[-(z - z')^2/w^2]$ and phenotype-specific birth rate $b_i(z) = 1 - z^2/\theta^2$ which is quadratic in nature ([(4)]. Since we assume non-consumptive interference as trait-independent, the interference factor becomes :

$$Q_i(\mathbf{N}) = \frac{1}{1 + c_i\Big(\sum_{j=1}^{S} \eta_{ij} N_j\Big)^{\gamma}} \tag{S36}$$

and thus carries no dependence on $z$ and is thus a species-level quantity. This is a useful simplification: trait averages factorise exactly, with no and no numerical quadrature required to solve these equations numerically. Integrating Eq. S35 against $p_i(z)$ and using standard Gaussian moments gives the species-level dynamics

$$\frac{\mathrm{d}N_i}{\mathrm{d}t} = N_i \left[ b_i Q_i - m_0 - Q_i \sum_{j=1}^{S} Q_j\, \alpha_{ij} N_j \right], \tag{S37}$$

where the intraspecific term is recovered as the special case $j = i$, and thus carries $Q_i^2$, while interspecific terms carry the product $Q_i Q_j$. The trait-averaged birth rate and competition coefficients are :

$$b_i = \int \left(1 - \frac{z^2}{\theta^2}\right) p_i(z)\, \mathrm{d}z = 1 - \frac{\mu_i^2 + \sigma_i^2}{\theta^2}, \tag{S38}$$

$$\alpha_{ij} = \iint p_i(z)\, \alpha(z, z')\, p_j(z')\, \mathrm{d}z' \mathrm{d}z = \frac{w}{\sqrt{w^2 + 2\sigma_i^2 + 2\sigma_j^2}} \exp\left(-\frac{(\mu_i - \mu_j)^2}{w^2 + 2\sigma_i^2 + 2\sigma_j^2}\right), \tag{S39}$$

following Pastore et al[(4)]. Note that $\alpha_{ij} = \alpha_{ji}$: the Gaussian kernel yields a symmetric competition matrix, a fact we use below. Writing $d = \mu_1 - \mu_2$ and $\Sigma = w^2 + 2\sigma_1^2 + 2\sigma_2^2$, the two-species coefficients are

$$\alpha_{11} = \frac{w}{\sqrt{w^2 + 4\sigma_1^2}}, \qquad \alpha_{22} = \frac{w}{\sqrt{w^2 + 4\sigma_2^2}}, \qquad \alpha_{12} = \alpha_{21} = \frac{w}{\sqrt{\Sigma}}\, \mathrm{e}^{-d^2/\Sigma}. \tag{S40}$$

##### 3.1 Niche overlap and fitness ration

Multiplying both mutual-invasibility inequalities by $\sqrt{\alpha_{22}\alpha_{21}/(\alpha_{11}\alpha_{12})}$ recovers the standard MCT decomposition[(5)]. The niche overlap

$$\rho = \sqrt{\frac{\alpha_{12}\alpha_{21}}{\alpha_{11}\alpha_{22}}} \tag{S41}$$

depends only on the competition matrix and is thus unaffected by interference. Substituting Eq. S40,

$$\rho = \exp\left(-\frac{(\mu_1 - \mu_2)^2}{w^2 + 2\sigma_1^2 + 2\sigma_2^2}\right) \left[\frac{(w^2 + 4\sigma_1^2)(w^2 + 4\sigma_2^2)}{(w^2 + 2\sigma_1^2 + 2\sigma_2^2)^2}\right]^{1/4} \tag{S42}$$

identical to the expression obtained for the Lotka-Volterra eco-evolutionary model[(4)]. When $\sigma_1 = \sigma_2 = \sigma$ this reduces to $\rho = \exp[-d^2/(w^2 + 4\sigma^2)]$: intraspecific trait variation inflates the effective kernel width and thereby raises niche overlap at any given trait separation.

Since the Gaussian kernel gives $\alpha_{12} = \alpha_{21}$ (Eq. S40), the Chesson factor simplifies to

$$\sqrt{\frac{\alpha_{22}\alpha_{21}}{\alpha_{11}\alpha_{12}}} = \sqrt{\frac{\alpha_{22}}{\alpha_{11}}} = \left[\frac{w^2 + 4\sigma_1^2}{w^2 + 4\sigma_2^2}\right]^{1/4}. \tag{S43}$$

The fitness ratio thus becomes

$$\left.\frac{\kappa_1}{\kappa_2}\right|^{(j)} = \frac{\tilde{R}_1^{(j)}}{\tilde{R}_2^{(j)}}\left[\frac{w^2+4\sigma_1^2}{w^2+4\sigma_2^2}\right]^{1/4}, \qquad j\in\{1,2\}. \tag{S44}$$

Unlike $\rho$, this quantity is resident-conditional: it takes a different value depending on which species is at equilibrium, Since $Q_i^{(1)} \neq Q_i^{(2)}$ whenever $\eta_{i1}K_1 \neq \eta_{i2}K_2$. Explicitly, using Eqs. S38 and S9,

$$\left.\frac{\kappa_1}{\kappa_2}\right|^{(j)} = \frac{1-m_0-\dfrac{\mu_1^2+\sigma_1^2}{\theta^2}-m_0c_1\big(\eta_{1j}K_j\big)^\gamma}{1-m_0-\dfrac{\mu_2^2+\sigma_2^2}{\theta^2}-m_0c_2\big(\eta_{2j}K_j\big)^\gamma}\left[\frac{w^2+4\sigma_1^2}{w^2+4\sigma_2^2}\right]^{1/4}. \tag{S45}$$

##### 3.2 Coexistence condition

Mutual invasibility requires both inequalities of Eq. S8, which in MCT coordinates read

$$\rho < \left.\frac{\kappa_1}{\kappa_2}\right|^{(2)} \qquad \text{and} \qquad \left.\frac{\kappa_1}{\kappa_2}\right|^{(1)} < \frac{1}{\rho} \tag{S46}$$

Both conditions must hold, and each constrains a different fitness ratio, evaluated at a different resident equilibrium.

##### 3.3 The Lotka-Volterra limit

Setting $c_i = 0$ (equivalently $\eta_{ij} = 0$) gives $Q_i \equiv 1$ and $\tilde{R}_i^{(j)} \to r_i = b_i - m_0$ irrespective of the resident's identity. The two fitness ratios then collapse to the single value

$$\frac{\kappa_1}{\kappa_2} = \frac{\theta^2(1-m_0)-\mu_1^2-\sigma_1^2}{\theta^2(1-m_0)-\mu_2^2-\sigma_2^2}\left[\frac{w^2+4\sigma_1^2}{w^2+4\sigma_2^2}\right]^{1/4}, \tag{S47}$$

recovering the classical sandwich $\rho < \kappa_1/\kappa_2 < 1/\rho$ and Eq. S40 as in Pastore et al[(4)]. If additionally $\sigma_1 = \sigma_2$, the bracketed term equals unity and the fitness ratio reduces to the ratio of trait-averaged birth rates.

##### 3.4 Stability Analysis

We compute and plot the region of stable coexistence for the MCT space of fitness difference and niche difference. In classical MCT, the region of feasible and stable coexistence overlaps absolutely for GLV, meaning stability is guaranteed if Chesson's feasibility criteria are fulfilled. Here, we show that stability is also guaranteed if feasibility criteria (S46) are fulfilled for the interference model. We derive the Jacobian for the two-species sublinear model (S23),

$$J_{1j} = N_1\left[b_1\frac{\partial Q_1}{\partial N_j} - \alpha_{1j}Q_1Q_j - 2\alpha_{11}Q_1\frac{\partial Q_1}{\partial N_j}N_1 - \alpha_{12}N_2\left(\frac{\partial Q_1Q_2}{\partial N_j}\right)\right] \tag{S48}$$

$$J_{2j} = N_2\left[b_2\frac{\partial Q_2}{\partial N_j} - \alpha_{2j}Q_2Q_j - 2\alpha_{22}Q_2\frac{\partial Q_2}{\partial N_j}N_2 - \alpha_{21}N_1\left(\frac{\partial Q_1Q_2}{\partial N_j}\right)\right] \tag{S49}$$

Where,

$$\frac{\partial Q_i}{\partial N_j} = -\gamma c_i\eta_{ij}Q_i^2\left(\sum_k \eta_{ik}N_k\right)^{\gamma-1} \tag{S50}$$

In Figure S1, the stable region is marked where the largest eigenvalue of $J$ is less than zero, and it absolutely overlaps with the feasibility region.

##### 3.5 Construction and standardisation of interference series

We kept only experiments in which consumer number, resource density and the dimensions of the experimental arena were reported and in which feeding could be expressed on a per-consumer basis. We converted observations to common density and time units within each series, while parameters were allowed to vary among series. Here in the experiments absolute space-clearance rates and handling times are not directly comparable among different taxa and foraging dimensions. We used consumer-free controls where available, to correct for background resource mortality. We excluded experiments from the primary analysis if consumer density covaried with temperature, consumer size, resource identity, habitat complexity or experimental duration, or if consumers could emigrate from the arena. We analysed two-dimensional and three-dimensional foragers using density measured per unit area and per unit volume, respectively. Systems classified as 2.5-dimensional in FoRAGE were excluded from the primary analysis since the appropriate density measure is not uniquely defined.

| Symbol | Description |
|---|---|
| $N_i$ | Population density of species $i$. Their initial value is 1 for all species. |
| $\mu_i$ | Mean trait value of species $i$. Initial values are sampled from $U[-0.7,\ 0.7]$ |
| $\sigma_i$ | Trait standard deviation of species $i$; sampled from either $U[0.001,\ 0.005]$ (low), or $U[0.01,\ 0.1]$ (high). |
| $h_i^2$ | Heritability of species $i$'s trait for two species and subsequent $S$ species; sampled from $U[0.1,\ 0.15]$ for $S$ species, but for two species it is $\in (0.1, 0.15)$ |
| $\theta$ | Width of intrinsic growth curve; 0.75. |
| $\omega$ | Width of competition kernel at 0.1. |
| $c_i$ | interference sensitivity fixed at 1 for all species |
| $\gamma$ | interference scaling exponent quantifying behavioural mediated interference, either 0.5 or 1 for nearest-neighbour interference or well-mixed respectively. |
| $m_0$ | intrinsic mortality rate fixed at 0.25 for all species. |
| $\eta_{ij}$ | interference coefficient of species $j$ on $i$, capturing non-consumptive interactions. |
| $\alpha_{ij}$ | competitive effect of species $j$ on $i$. |
| $\beta_{ij}$ | selective competitive pressure of species $j$ on species $i$. |

Table S1: Parameters and their values with description used in the study for the dynamical simulations. $U[a,\ b]$ is the uniform distribution between $a$ and $b$.

| Symbol | IBM parameter | Description |
|---|---|---|
| $N$ | `N` | Population abundance; variable. |
| $\xi$ | `xi` | Individual behavioural parameter. For $\xi > 0$, individuals are likely to interfere with or move towards their nearest neighbours (e.g., aggregation); for $\xi < 0$, individuals are likely to move away from or repel from their local neighbourhood; and $\xi = 0$ corresponds to movement in random directions. $\xi \in \{-1, -0.75, -0.5, 0, 0.5, 0.75, 1\}$. |
| $R$ | `arena_radius` | Radius of the circular arena, set to 10.0. |
| $r$ | `larva_radius` | Radius of an individual larva, set to 0.45. |
| $f$ | `feed_rate` | Individual feeding rate, set to 0.1. |
| $v$ | `move_speed` | Individual movement speed, drawn from $\mathcal{N}(0.5, 0.5)$. |
| $\tau_p$ | `penalty_time` | Number of time steps penalised following interference or collision, set to 8.0. For the LV model, the penalty is set to 0.0. |
| $m_0$ | `m0` | Intrinsic mortality energy threshold, set to 5.0. |
| $\epsilon$ | `conversion_efficiency` | Conversion efficiency of consumed resources into offspring, set to 0.2. |
| $R_0$ | `total_resource` | Total amount of resources initially available to the population, set to 2500. |
| $r_{\text{int}}$ | `r_int` | Interaction radius for attraction or repulsion, set to 1.0. |
| $T_{\max}$ | `t_max` | Total duration of the foraging simulation, set to 500 time steps. |

Table S2: Parameters used in the individual-based model (IBM). IBM parameter names are shown in typewriter font to match their names in the model implementation. $\mathcal{N}(a, b)$ denotes a normal distribution with mean $a$ and standard deviation $b$.

### References


[1] Arthur, R. M. Species packing, and what competition minimizes. *Proceedings of the National Academy of Sciences* **64**, 1369–1371 (1969).

[2] Baruah, G., Barabás, G. & John, R. When Do Trait-Based Higher Order Interactions and Individual Variation Promote Robust Species Coexistence? *Ecology and Evolution* **15**, e71336 (2025). URL https://onlinelibrary.wiley.com/doi/abs/10.1002/ece3.71336. _eprint: https://onlinelibrary.wiley.com/doi/pdf/10.1002/ece3.71336.

[3] Barabas, G. & D'Andrea, R. The effect of intraspecific variation and heritability on community pattern and robustness. *Ecology Letters* **19**, 977–986 (2016).

[4] Pastore, A. I., Barabás, G., Bimler, M. D., Mayfield, M. M. & Miller, T. E. The evolution of niche overlap and competitive differences. *Nature Ecology & Evolution* **5**, 330–337 (2021). URL https://www.nature.com/articles/s41559-020-01383-y. Number: 3 Publisher: Nature Publishing Group.

[5] Chesson, P. Mechanisms of Maintenance of Species Diversity. *Annual Review of Ecology and Systematics* **31**, 343–366 (2000). URL http://www.annualreviews.org/doi/10.1146/annurev.ecolsys.31.1.343.

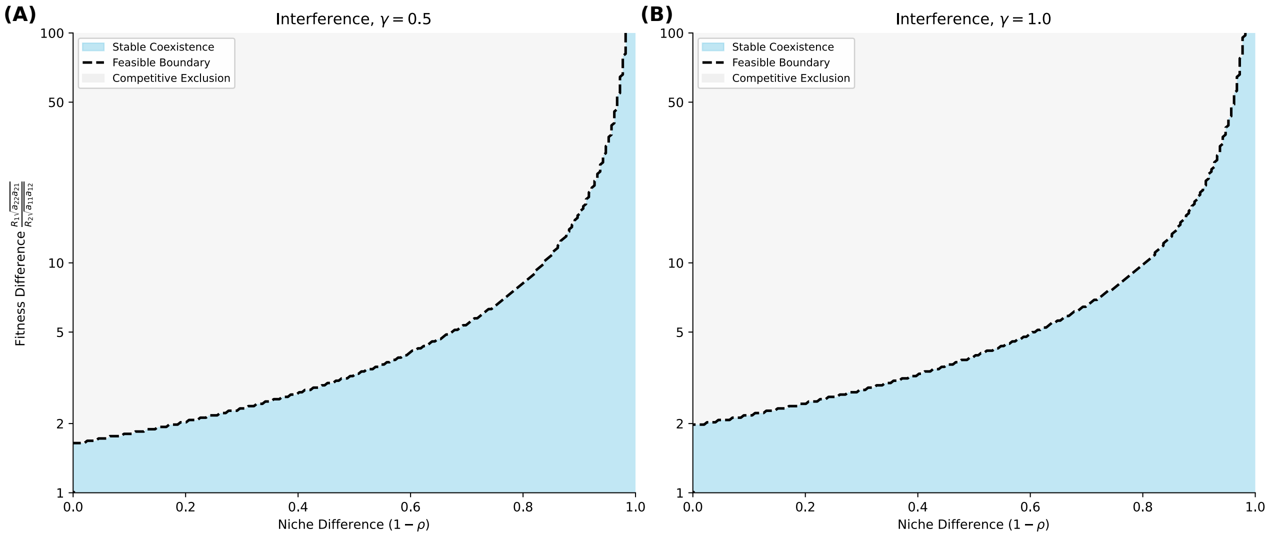


Figure S1: **Feasibility guarantees stability in the interference model.** Stable Coexistence regions in the MCT space of fitness ratio($\frac{R_1}{R_2}\sqrt{\frac{\alpha_{22}\alpha_{21}}{\alpha_{11}\alpha_{12}}}$) and niche difference $(1-\rho)$. Blue color indicates parameter combinations satisfying mutual invasibility, and the corresponding largest eigenvalue of the Jacobian is a negative real number. Grey color indicates competitive exclusion. **(A)** $\gamma = 0.5$ **(B)** $\gamma = 1.0$ . Parameter values $b_i = 1.0, \alpha_{11} = \alpha_{22} = 1, m_0 = 0.25, c_i = 1, \eta_{11} = \eta_{22} = 1.0, \eta_{12} = \eta_{21} = 0.2$

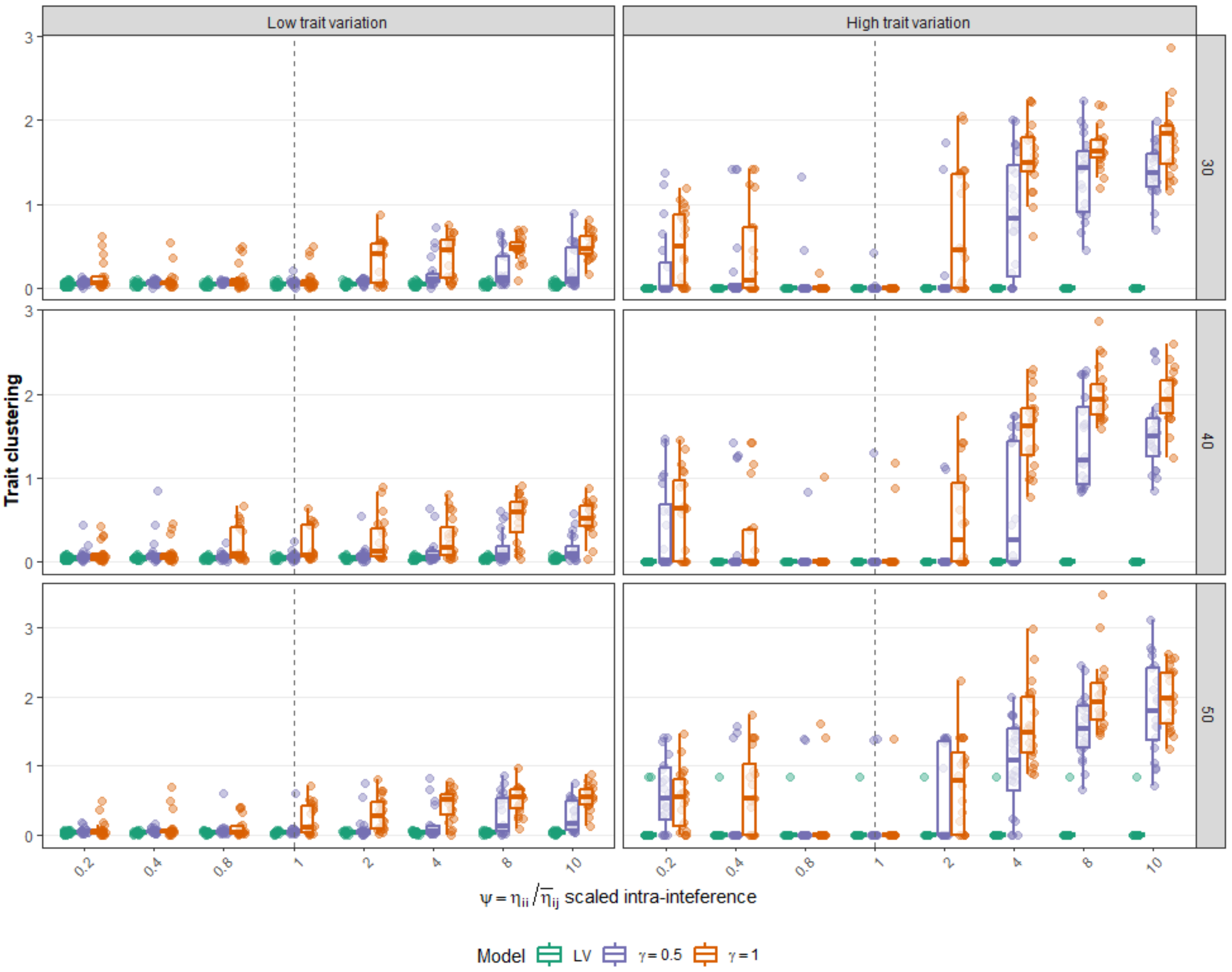


Figure S2: **Species trait clustering in larger competitive communities.** For initial $S = 30, 40, 50$ species competitive community (row panels), in eco-evolutionary equilibria species trait clustering (coefficient of variation in equilibrium trait values) was plotted against the ratio of intra-interference versus inter-interference factor, i.e., $\psi = \frac{\eta_{ii}}{\eta_{ij}}$, for sublinear models with $\gamma = 0.5$ (violet color boxplots), 1 (orange color boxplots), and generalised Lotka-Volterra model (green boxplots). $\gamma = 0.5$ represents nearest-neigbour interference mechanism, and $\gamma = 1$ represents well-mixed interference mechanism. High values of trait clustering indicates high trait convergence. In general, with sublinear models, clustering is generally tolerated. Boxplots consists of 40 independent replicate simulations of our eco-evolutionary model (equation 21, 22). Parameters include from table S1.

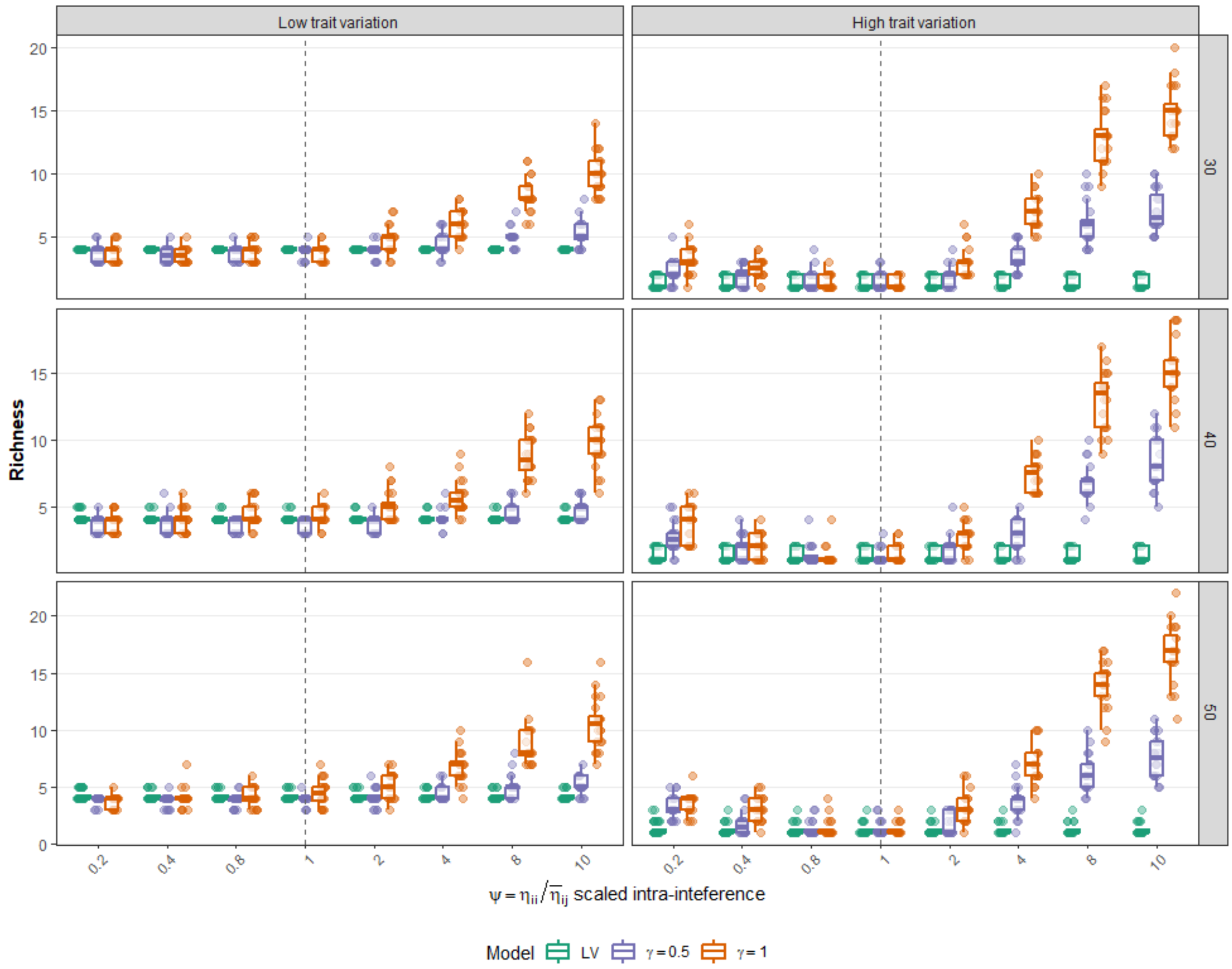


Figure S3: **Species coexistence in larger competitive communities in sublinear density regime.** For initial $S = 30, 40, 50$ species competitive community (row panels), in eco-evolutionary equilibria species richness ( total species above an extinction threshold of density $10^{-6}$) was plotted against the ratio of intra-interference versus inter-interference factor, i.e., $\psi = \frac{\eta_{ii}}{\eta_{ij}}$, for sublinear models with $\gamma = 0.5$ (violet color boxplots), 1 (orange color boxplots), and generalised Lotka-Volterra model (green boxplots). $\gamma = 0.5$ represents nearest-neigbour interference mechanism, and $\gamma = 1$ represents well-mixed interference mechanism. For generalised trait-based LV model, high trait variance (right figure column) leads to lower amount of species coexisting across different initial species community in contrast to low trait variation and sublinear models. Boxplots consists of 40 independent replicate simulations of our eco-evolutionary model (equation 21, 22). Parameters include from table S1.

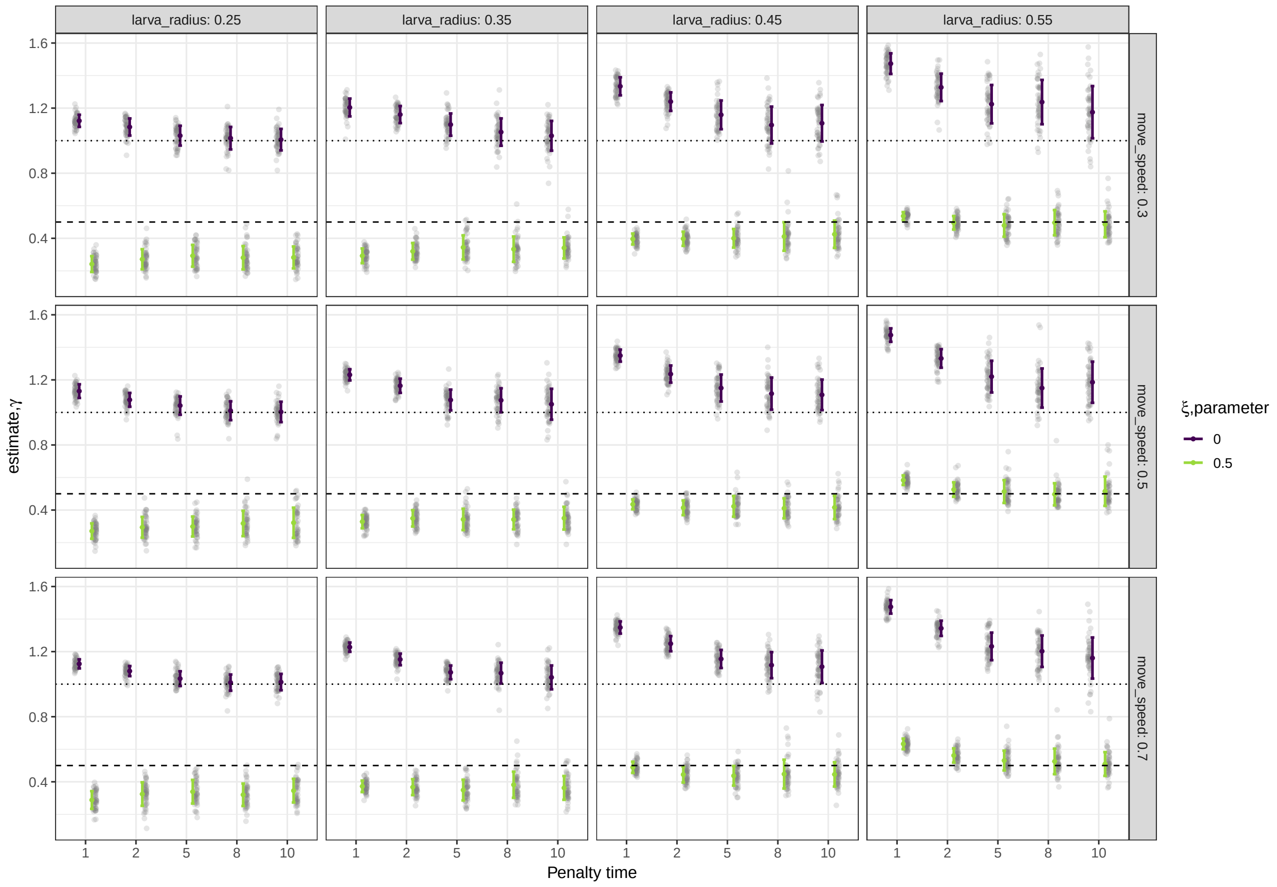


Figure S4: **Sensitivity simulations on estimated interference scaling factor $\gamma$ for different parameters of the IBM**. For $\xi$ parameter of 0 and 0.5 signifying well-mixed random movement behaviour, and nearest-neighbour mechanism respectively. Under different consumer size (`larva_radius`), and penalty time for interference (x-axis), and different consumer speeds (`move_speed`), we observe some variation in the estimation of $\gamma$ under two different $\xi$ mechanisms. Grey dots indicate replicate simulations (n=50) with mean and standard deviations. Horizontal dotted black line and horizontal dashed black line represent the estimate of the interference scaling factor $\gamma = 1$ and $\gamma = 0.5$ from theory, Box 1 and Box 2.
